# Cochlear transcriptome analysis of an outbred mouse population (CFW)

**DOI:** 10.1101/2023.02.15.528661

**Authors:** Ely Cheikh Boussaty, Neil Tedeschi, Mark Novotny, Yuzuru Ninoyu, Eric Du, Clara Draf, Yun Zhang, Uri Manor, Richard H. Scheuermann, Rick Friedman

## Abstract

Age-related hearing loss (ARHL) is the most common cause of hearing loss and one of the most prevalent conditions affecting the elderly worldwide. Despite evidence from our lab and others about its polygenic nature, little is known about the specific genes, cell types and pathways involved in ARHL, impeding the development of therapeutic interventions. In this manuscript, we describe, for the first time, the complete cell-type specific transcriptome of the aging mouse cochlea using snRNA-seq in an outbred mouse model in relation to auditory threshold variation. Cochlear cell types were identified using unsupervised clustering and annotated via a three-tiered approach - first by linking to expression of known marker genes, then using the NS-Forest algorithm to select minimum cluster-specific marker genes and reduce dimensional feature space for statistical comparison of our clusters with existing publicly-available data sets on the gEAR website (https://umgear.org/), and finally, by validating and refining the annotations using Multiplexed Error Robust Fluorescence In Situ Hybridization (MERFISH) and the cluster-specific marker genes as probes. We report on 60 unique cell-types expanding the number of defined cochlear cell types by more than two times. Importantly, we show significant specific cell type increases and decreases associated with loss of hearing acuity implicating specific subsets of hair cell subtypes, ganglion cell subtypes, and cell subtypes withing the stria vascularis in this model of ARHL. These results provide a view into the cellular and molecular mechanisms responsible for age-related hearing loss and pathways for therapeutic targeting.

## Introduction

Progressive, bilateral sensorineural hearing impairment affects approximately 25% of people aged 65-74 and 50% aged 75 and older. In the United States, two thirds of people over 70 years of age have some degree of hearing loss (Bainbridge and Wallhagen, 2014; Gurgel et al., 2014; Jayakody et al., 2018). Aside from the detrimental impact on quality of life, hearing loss also carries an increasing economic burden as the cost of medical expenditures is expected to reach $60 billion in 2030. It is projected that 2.45 billion people will have hearing loss by 2050, a 56.1% increase from 2019, despite stable age-standardized prevalence (Vos et al., 2020). Notably, a substantial fraction of patients with progressive hearing loss have no identifiable mutation in any known hearing loss gene among the 100+ that have been identified, suggesting that a significant fraction of hearing loss is due to unidentified monogenic or polygenic causes (Bowl and Dawson, 2019).

As part of a broad approach to studying the genetic landscape of ARHL, we have begun a large-scale effort to phenotypically characterize the auditory function of young and aged Carworth Farms White (CFW) Crl:CFW(SW)-US_P08 (hereafter CFW) outbred mice (Parker et al., 2016). Although not specifically developed for genetic research, these mice have several attractive properties for gene discovery. CFW mice were derived from a small number of founders and have been maintained as an outbred population for more than 100 generations, thus, reducing the size of linkage disequilibrium between alleles (Rice and O’Brien).

Developing therapies for progressive hearing loss necessitates an understanding of the genes and pathways involved. The cellular complexity of the inner ear, including the sensory and supporting cells of the organ of Corti, the lateral wall (stria vascularis) and the auditory neurons, necessitates a single cell approach to gain an understanding of the pathway changes associated with hearing loss. Due to this complexity, a few laboratories have used single-cell RNA sequencing (scRNA-seq) to characterize the molecular mechanisms underlying cochlear development^14^ and to gain insights into aging in a single mouse strain, and after acoustic trauma (Hoa et al., 2020; Kolla t al., 2020; Korrapati et al., 2019; Petitpré et al., 2022; Shrestha et al., 2018; Sun et al. 2018; Petitpréet al. 2022) . This approach has provided insights into the molecular mechanisms underlying cochlear development (Brown et al., 2008) and response to damaging noise (Lavinsky et al., 2015; Liu et al., 2021). Recently, Li et al. looked at the transcriptome of inner and outer hair cells and the stria vascularis in aging CBA/J mice, implicating changes in several processes including gene transcription, DNA damage, autophagy and oxidative stress (Li et al., 2018). However, these single cell analyses of the adult mouse inner ear, particularly the organ of Corti, have been hampered by the difficulty in dissociating cells from the tissue due to their tight junction connections and their extracellular matrix embeddings (Burns et al., 2015).

In this study, we used *single nucleus RNA-seq (snRNA-seq)* across 48 genetically diverse CFW outbred mice at 10 months of age to provide for a more unbiased representation of cell types associated with variations in hearing. We identified gene expression signatures for 60 distinct cell types withing the cochlea, including novel markers for inner and outer hair cell subtypes and found that specific hair cell, ganglion, and stria vascularis cell types show preferential depletion associated with hearing loss. To our knowledge this is the first snRNA-seq study to examine differential gene expression across all cochlear cell types in genetically diverse outbred mice with varying degrees of hearing loss.

## Results

CFW mice are a genetically diverse outbred mouse population distinguished by degrading linkage disequilibrium (LD) between nearby alleles and shorter LD ranges compared to other commercially available inbred mouse strains (Rice and O’Brien, 1980; Chia et al., 2005), which makes them ideal models to capture high-resolution mapping in genome-wide association studies (GWAS). Our ongoing efforts to phenotype the hearing function in CFW mice have categorized them into eight distinct patterns of hearing, ranging from normal hearing, to moderate mid- or high-frequency hearing loss, to profound hearing loss at all frequencies, with each of those hearing patterns worsening with age (Du et al 2022).

A group of 48 mice were selected to represent those hearing patterns (Figure 1), based on their proportion in the general CFW cohort, to explore the molecular and cellular correlates of hearing loss using single nucleus RNA sequencing (snRNA-seq) of dissected cochlea. During the dissection of inner ear tissue, special care was given to quickly process the samples while avoiding any shear force. The vestibular parts of the inner ear, including utricle, saccule, and semicircular canal ampulla, were removed before collecting cochlear tissue. snRNA-seq processing was chosen over scRNA-seq to avoid the stress responses induced during the single cell dissociation procedure and to provide for a more unbiased representation of cell types from solid tissues (Bakken et al., 2018).

**Figure 1.**
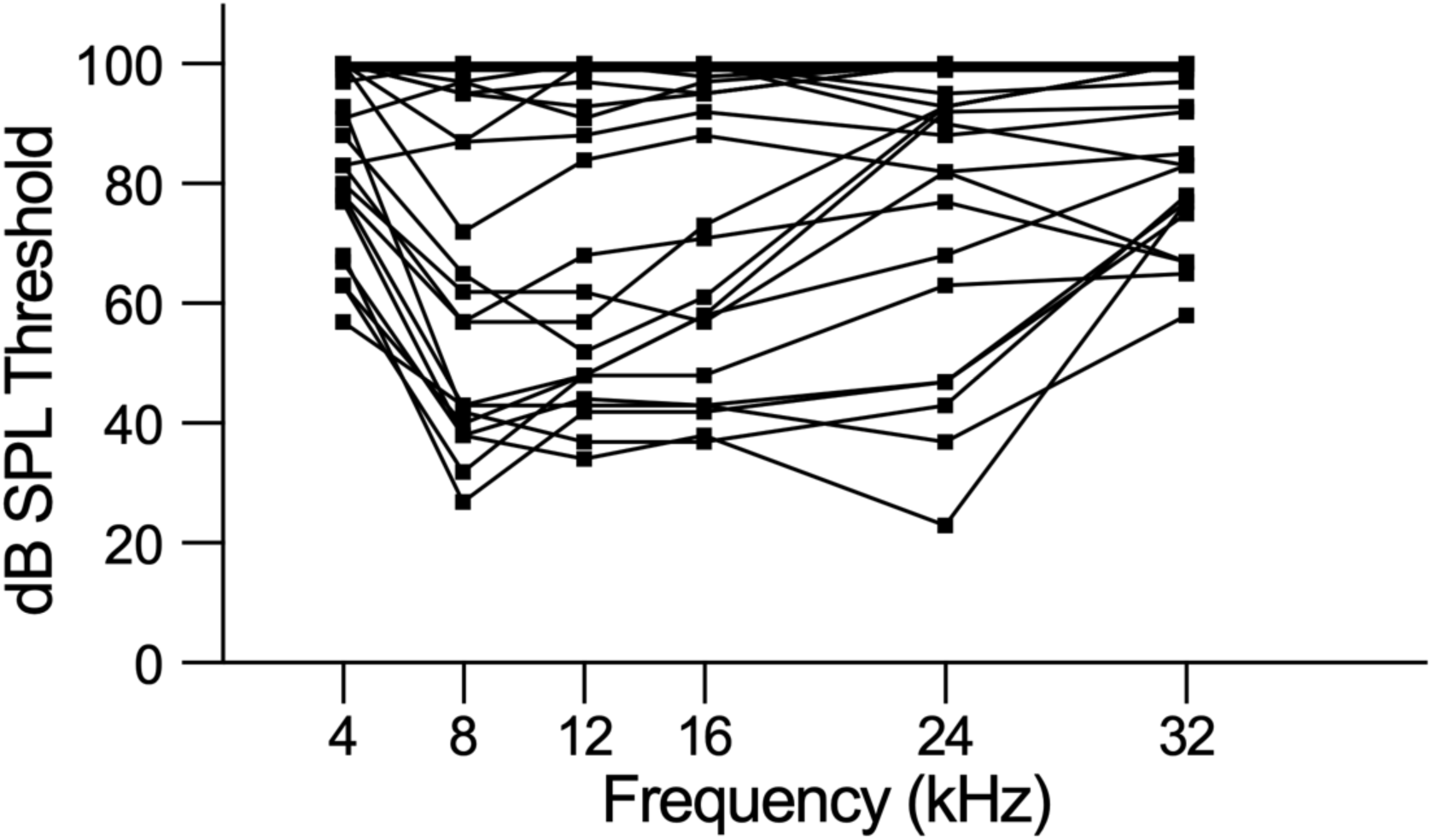
– Phenotyping CFW outbred mice shows eight distinct hearing patterns. Individual Auditory Brain Response (ABR) thresholds plots for mice selected to represent CFW cohort hearing patterns for cochlear tissue collection and snRNA-seq analysis.

Iterative unsupervised clustering was used to group snRNA-seq transcriptional profiles into transcriptome clusters. The initial Leiden community detection yielded 26 clusters, a subset of which (13) were processed through a second round of sub-clustering based on Silhouette score and manual inspection to yield a final collection of 60 transcriptome clusters (Figure 2A). The similarity relationships between the clusters were determined using hierarchical clustering (Figure 2B) and the relative abundance of each putative cell type determined (Figure 2C). The hierarchical clustering results confirmed the transcriptional relationships between the initial clusters and the sub-clusters with common origins from the second round of clustering.

**Figure 2.**
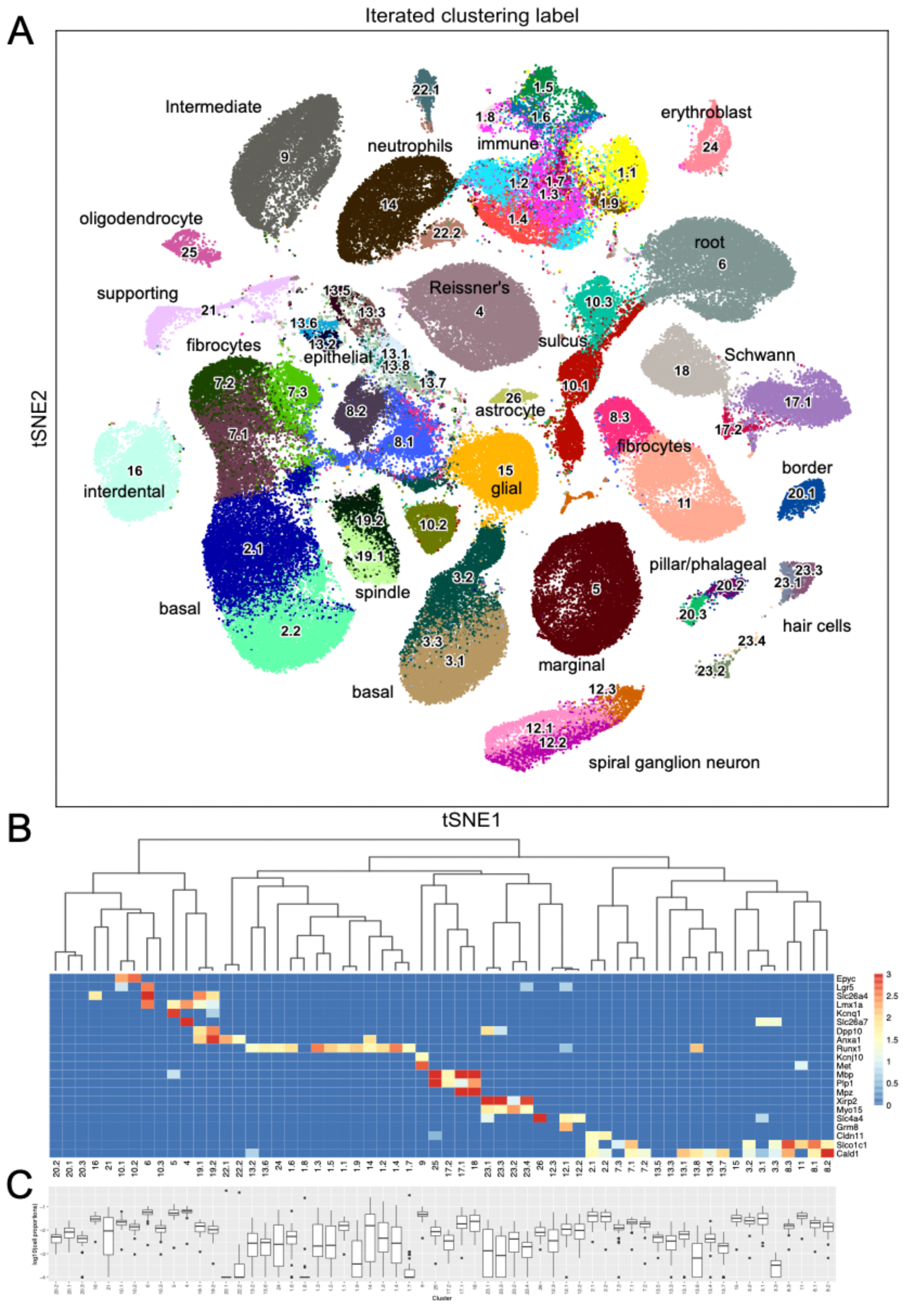
– Sixty snRNA-seq transcriptomic clusters from mouse cochlea. A) Iterative unsupervised clustering of the processed cell-by-gene snRNA-seq expression matrix performed using Leiden community detection identified sixty distinct transcriptomic clusters (numbers). tSNE embedding of the clustering results is shown. Cell type annotations of each transcriptomic cluster based on subsequent analysis (Table 1) is shown. The data can be explored at https://umgear.org/index.html?multigene_plots=0&layout_id=c2acc279&gene_symbol_exact_match=1&gene_symbol=slc26a5. B) Heatmap of marker genes known to be expressed in specific ear cell types arranged according to hierarchical clustering of transcriptomic clusters, which captures the transcriptional relationships between the discrete clusters. The horizontal banding patterns of these known marker genes validates the structure of the hierarchical clustering taxonomy. C) Box plots of the proportion (log10 transformed) of each cell type across the 48 outbred mouse cochlear samples, with median proportion ranging from ∼10^-4^ for Cluster 22.1 to ∼10^-1^ for Cluster 4.

As an initial approach toward determining which cell types correspond to these transcriptomic clusters, we examined the expression levels of several genes known to be expressed in specific ear cell types (Figure 2B). The organ of Corti supporting cell gene Epyc (Hanada et al., 2017) is uniquely expressed in the two related Clusters 10.1 and 10.2. The inner ear progenitor gene Lgr5 (Chen et al., 2021) is uniquely expressed in Cluster 6. The spindle-root gene Slc26a4 is uniquely expressed in the two related Clusters 19.1 and 19.2 (Nishio et al., 2016) . The Kcnq1 marginal cell gene and the Slc26a7 Reissner’s membrane gene are preferentially expressed in Cluster 5 and Cluster 4, respectively. The hematopoietic stem cell-related gene Runx1 (de Bruijn & Dzierzak, 2017) is expressed in twelve related clusters. The intermediate cell genes Kcnj10 (Marcus et al., 2002) and Met (Shibata et al., 2016) are uniquely and preferentially expressed in Cluster 9. The glial and Schwann’s cell genes Mpz, Mbp, and Plp1 are expressed in the related Clusters 25, 17.1, 17.2, and 18. The hair cell (HC) genes Xirp2 (Scheffer et al., 2015) and Cabp2 (Yang et al., 2016) are expressed in related Clusters 23.1, 23.2, 23.3, and 23.4. The spiral ganglion neuron (SGN) genes Slc4a4 (Grandi et al., 2020) and Grm8 (Sun et al., 2021) are expressed in related Clusters 12.1 and 12.2. The basal cell gene Cldn11 (Gow et al., 2004) is expressed in related Clusters 2.1 and 2.2. Finally, the supporting fibrocyte genes Slco1c1 (Ng et al., 2021) and Cald1 (Scheffer et al., 2015) are expressed in seventeen related clusters. While the expression pattern of known marker genes was useful in providing an initial annotation and validation of the snRNA-seq analysis, there were many examples where finer grained cell type distinctions were identified from the unsupervised clustering results.

Therefore, to further extend this cell type classification, the NS-Forest algorithm was used to identify the minimum sets of marker genes for each cluster. NS-Forest uses random forest machine learning and a binary expression scoring method to identify necessary and sufficient marker genes, optimally capturing the essence of cell type identity (Aevermann et al., 2018; 2021). NS-Forest analysis of the 60 cell type clusters yielded 117 marker genes with high cell type specificity as illustrated by the diagonal expression pattern of markers across the clustered dataset (Figure 3A) and the relatively high F-beta values of classification accuracy (Supplementary Table 1). For example, HC Clusters 23.2 and 23.4 showed F-beta values of 0.93 and 0.88 using the single cell type markers Slc26A5 (Yamashita et al., 2015) and Ripor3, respectively. For the other hair cell clusters, combinatorial expression of two marker genes gave optimal classification accuracy of F-beta = 0.85 with Gm1113 and Ofcc1 markers for Clusters 23.1 and markers Ush2a and C230072F16Rik for Cluster 23.3. NS-Forest marker genes for the SGN populations included Ntng1, Mdga1, Cdh9, and Meg3, which have been shown to be among the highest differentially expressed genes during the diversification process of SGN (Petitpré et al., 2022).

**Figure 3.**
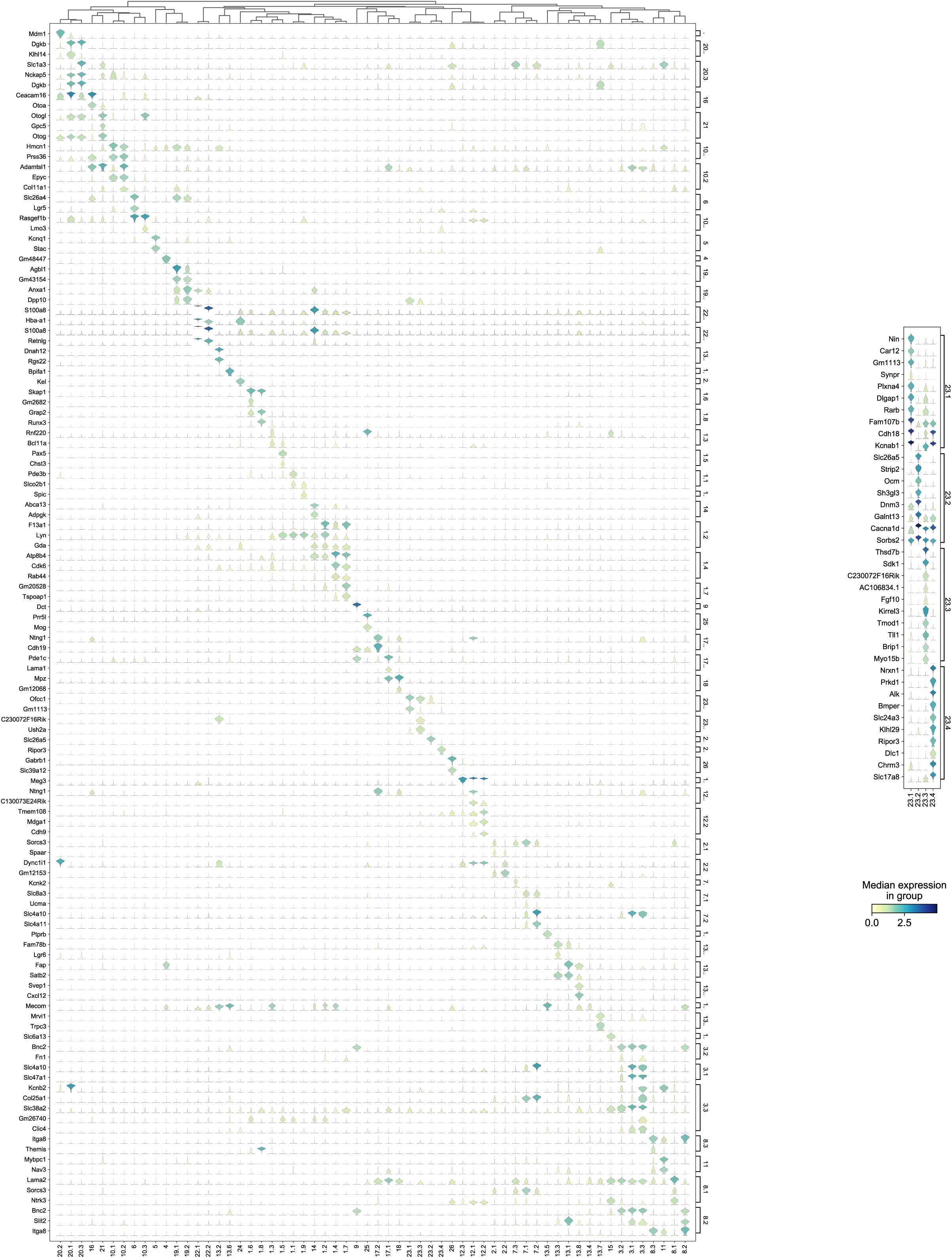
– Cell type-specific marker genes. NS-Forest was used to identify the minimum set of necessary and specific marker genes for each cell type, with gene expression (rows) across each transcriptomic cluster (columns) illustrated in violin plots. A) Minimum NS-Forest marker genes identified from the complete cell-by-gene expression matrix, with diagonal expression patterns and little off-diagonal expression showing marker gene specificity. In addition to the minimum site of marker genes, NS-Forest also produces an extended set of ten genes selected based on binary expression (on-off) patterns in the target cluster. B) Top ten binary genes were identified within the four Cluster 23 iterative subclusters using the pooled Cluster 23 subcluster cell-by-gene expression matrix as input, showing strong and distinct binary expression patterns, especially for Clusters 23.3 and 23.4.

NS-Forest also produces an extended set of marker genes that show specific binary expression patterns across the cell clusters. In the case of the four HC clusters (Clusters 23.1, 23.3, 23.2 and 23.4), the genes Slc26a5, Ocm, and Cacnald encoding for Prestin (Yamashita et al., 2015), Oncomodulin (Sakaguchi et al., 1998), and Calcium voltage dependent channel (Cav1.3) (Chen et al., 2012) respectively, are highly expressed in Cluster 23.2 suggesting this group is an outer hair cell (OHC) population (Figure 3B). Cluster 23.4 shows specific expression of Slc17a8 (Ruel et al., 2008) and Nrxn1 (Mozhui et al., 2011), known inner hair cell markers (IHC) encoding the Vesicular Glutamate Transporter-3 and a presynaptic adhesion protein, respectively. Cluster 23.3 shows specific expression of Myo15b, while Cluster 23.1 shows high expression of Kcnab1, which encodes a potassium voltage-gated channel, a hair cells marker conserved across species (Janesick et al., 2021) and shown to be more IHC specific in mature mouse cochlea (Liu et al., 2014; Bi et al., 2022). Genes recently validated as HC gene markers in newborn cochlea (Kolla et al., 2020) were not identified in our dataset, suggesting that those markers may be more specific for early postnatal stages.

To extend these granular cell type annotations further, the transcriptomics clusters were compared against five recently released snRNA and scRNA postnatal dataset (Supplementary Table 2) hosted by the gEAR database (https://umgear.org) using FR-Match, a cluster-to-cluster cell type matching algorithm that incorporates shared information among cells to determine the relationship between two transcriptome clusters (Zhang et al., 2021) using the cluster-specific marker genes identified by NS-Forest to provide a reduced dimensional feature space and support their statistical comparison. A match was considered high confidence if a match was found in both directions (query to reference and reference to query) and/or if a match was found to the same cell type reported in two or more gEAR datasets; a match was considered low confidence if there was a match in only one dataset in only one direction. The FR-Match comparisons show that 29 clusters were matched to gEAR cell types with high confidence (Supplementary Figure 1 and Table 1). The analysis results confirmed the classification of strial cells (marginal, intermediate, and basal) and spindle-root populations with high confidence (up to 8 matches). The clusters corresponding to cochlear outer hair cells, glial/Schwann cells, immune cells, and a subset of fibrocyte populations were also confirmed with high confidence, albeit with lower numbers of matches. The cell populations characterized as Hensen’s cells, Dieters cells, and pillar cells and the remaining fibrocytes populations were matched with lower confidence. The results of FR-Match analysis were also consistent with the cluster hierarchy, highlighting the advantage of using the FR-Match to identify corresponding cell types.

To validate the specificity of NS-Forest cell-type-specific marker expression and to characterize the spatial relationships between the cell types, the recently developed multiplexed error-robust fluorescent in situ hybridization technique (MERFISH), a single-molecule imaging approach that allows identification and spatial localizations of RNA transcripts for several different cell types (Moffitt and Zhuang 2016; Zhuang, 2021), was used in cross-sections from P5 cochleae. The NS-Forest cell-type specific marker genes (Figure 3A) were utilized as MERFISH probes to confirm the consistency between the cell-type annotations and transcript localizations. For example, marker genes for hair cell clusters localized to inner hair cells (IHC) (Cluster 23.1, Gm1113+/Ofcc1+) (Fig. 4A and C), outer hair cells (OHC) (Cluster 23.2, Slc26A5+) (Fig. 4B and C). Moreover, the MERFISH imaging confirmed the annotation of some subcluster categories with the lower confidence FR-Match results, such as border cells for Cluster 20.1 (Dgkb+/Klhl14+) (Fig. 4D), inner and outer pillar cells for Cluster 20.2 (Mdm1+) (Fig. 4E), Deiter’s cell and inner phalangeal cell for Cluster 20.3 (Dgkb+/Nckap5+/Slc1a3+) (Fig. 4F), where the localization pattern and clustering matched conventional anatomical localization nomenclature in the cochlea. Interestingly, some marker genes localized to the same anatomical cell types but indicated different cell populations among the subcategories, such as Cluster 12 for spiral ganglion cells (Fig. 4G and H) and Cluster 17 for Schwann cells (Fig. 4I). Cluster 12.2 cells strongly expressed Calbindin2 (Calb2) known for a marker of type1a spiral ganglion neuron (Shuohao 2018). These data suggested that the NS-Forest algorithm successfully identified unique marker genes for the transcriptomic clusters and the spatial transcriptomic analysis verified the FR-match annotations based on anatomical and physiological properties of cochlear cells.

**Figure 4.**
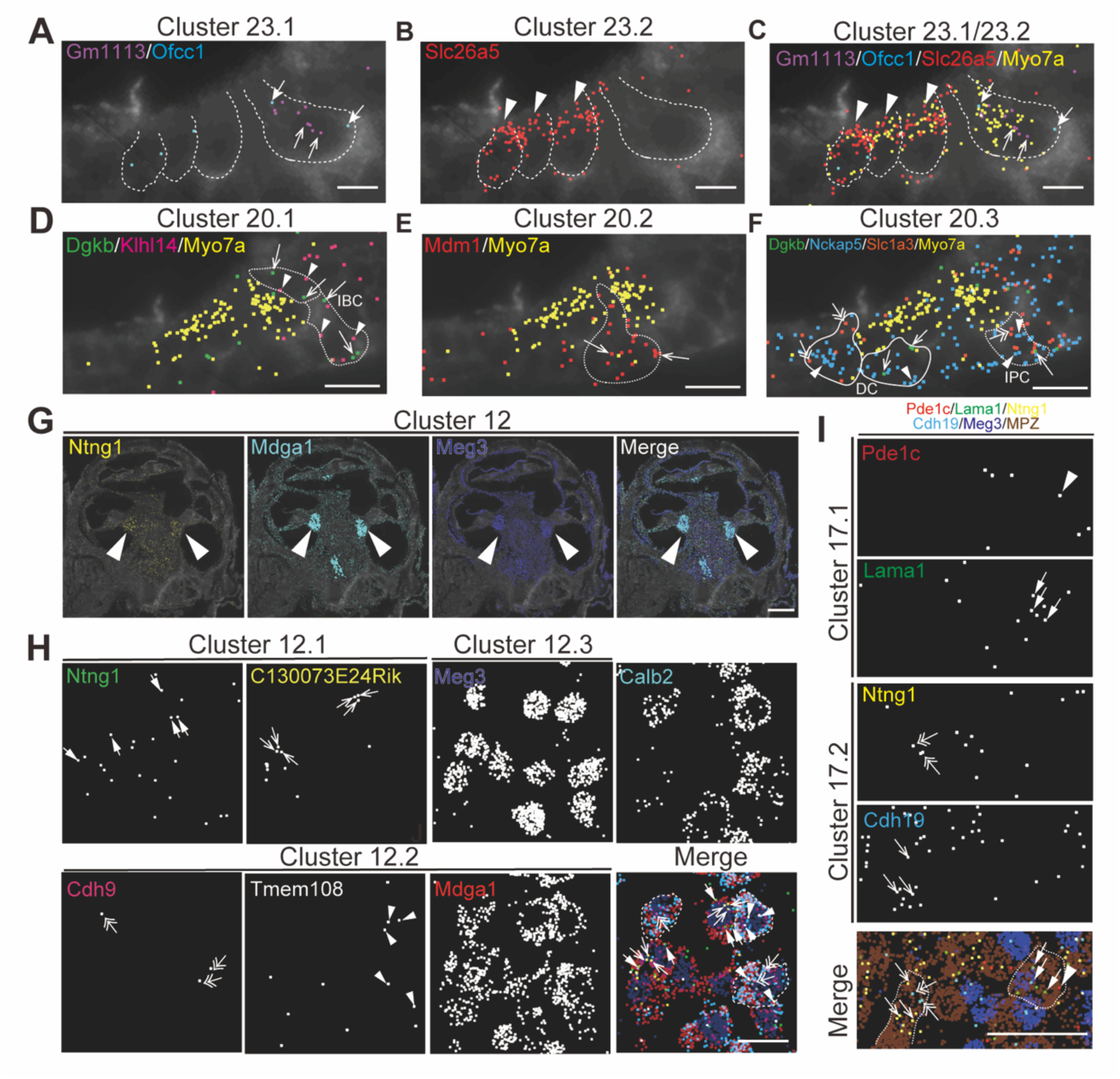
– Spatial organization of cell types in the mouse cochlea. MERFISH images with selected marker genes for Cluster 23 (cluster of hair cell) (A-C), Cluster 20 (cluster of supporting cell) (D-F), Cluster 12 (cluster of spiral ganglion cell) (G-H) and Cluster 17 (cluster of Schwann cell) (I) on the cochlear mid-modiolar cryosections at P4. Each pseudo-color dots represent transcripts of different marker genes and are merged with cell-boundary staining in dark grey. Cell boundaries are depicted by white or white-dashed line. Myo7a was used for pan-hair cell marker gene (yellow) (C-F). **A)** Cluster 23.1 for inner hair cell (IHC) with Gm113(purple, arrow), Ofcc1(light blue, thick arrow). **B)** Cluster 23.2 for outer hair cell (OHC) with Slc26a4 (red, arrow head). **C)** Cluster 23.4 for unknown hair-cell subtype with Ripor3 (white, arrow). **D)** Cluster 20.1 for inner border cell (IBC) with Dgkb (green, arrow) and Klhl14 (magenta, arrowhead). **E)** Cluster 20.2 for pillar cell with Mdm1 (red, arrow). **F)** Cluster 20.3 for inner phalangeal cell (IPC), Deiter’s cell (DC) and IBC with Dgkb (green, arrow), Nckap5 (blue, arrow head) and Slc1a3(orange, double arrow). **G)** Localization of marker genes for cluster 12 in cochlear spiral ganglion cells (arrow) with Ntng1 (yellow), Mdga1 (light blue) and Meg3 (dark blue). **H)** High magnification view of cluster 12.1 with Ntng1(green, thick arrow), C130073E24Rik (yellow, arrow), of cluster 12.2 with Cdh9 (magenta, double-head arrow), Tmem108 (white, arrowhead), and Mdga1 (red), and of cluster 12.3 with Meg3 (dark blue) and merged image with pseudo colors. Calbindin2 (calb2) is a known marker gene for type1a spiral ganglion neuron. Representative cell boundaries for Cluster 12.2 was depicted by white-dashed line. **M)** Images for cluster 17.1 with Pde1c (red, arrowhead) and Lama1(green, thick arrow), and for cluster 17.2 with Ntng1 (yellow, double-head arrow) and Cdh19 (light blue, arrow). Meg3 (dark blue) and Mpz (brown) indicate spiral ganglion cells and Schwann cells, respectively. Sale bars; 15μm (A-F), 250μm (G), and 25μm (H-J).

For further evaluation of the proportion of cell types present, we investigated the spatial organization of the scala media containing the organ of Corti, which is the receptor organ for hearing, by MERFISH (Supplementary Fig.2A and 2B). P5 aged mice were used to due to the difficulties in obtaining sufficient material in adult mice and to generate an atlas of cell types unbiased by age. In addition to hair cell and supporting cell clusters (Fig.4), we successfully identified various cell clusters in the cochlear duct, such as epithelial cells of Reisner membrane (Cluster 4, Soc26a7+/Tmem72+) (Supplementary Fig.2C), marginal cells (Cluster 5, Kcnq1+/Stac+) and intermediate cells (Cluster 9, Dct+) in the stria vascularis (Supplementary Fig.2D), spindle cell (Cluster 19.2, Anxa1+/Dpp10+) (Supplementary Fig.2E), root cells in the spiral prominence (Cluster 6, Lgr5+/Slc26a4+) (Supplementary Fig.2F), inner and outer sulcus cells (Cluster 10, Epyc+/Gata3+) (Supplementary Fig.2G), fibrocyte in the tympanic covering layer of basilar membrane (Cluster 11, Mybpc1+/Nav3) (Supplementary Fig.2H), and interdental cells in the spiral limbus (Cluster 16, Otoa+) (Supplementary Fig.2I). Notably, the clustering successfully distinguished transcriptional profiles even among rare cell types in the spiral prominence of the cochlea, including spindle cells (Cluster 19) and root cells (Cluster 6). Moreover, these clusters in the neonatal mouse cochlea were matched and confirmed with the adult mouse cochlea (Supplementary fig.3). These data demonstrate that the snRNA-seq approach robustly profiled cell type-specific transcriptomes in the cochlear tissue.

By combining the information derived from manual marker gene evaluation, gEAR reference matching, and MERFISH localization, cell type names and definitions for the 60 transcriptomics clusters were determined (Table 1). In order to provide for more descriptive cell types names for each of the transcriptomic clusters, we adopted a convention established by the ontology development community for defining and naming cell types derived from single cell transcriptomics experiments for incorporation into the Provisional Cell Ontology (Bakken et al., 2018; Tan et al., 2021), incorporating information about anatomic location, parent cell class, and specific marker gene combinations to provide for unique cell type names and experimentally useful definitions (Table 1 and Supplementary Table 1) to serve as future references.

To evaluate the cellular correlates of the loss of hearing acuity in this outbred population, the auditory brainstem response (ABR) thresholds were correlated with the cell type proportions in the 48 outbred CFW mice with three patterns observed (Figure 5A). For example, the proportion of cells in Clusters 23.2, 23.4, 12.1, and 12.2 showed an inverse correlation with ABR threshold (Figure 5A, red; Figure 5B, C, D), whereas the proportion of cells in Clusters 2.1 and 2.2 showed a positive correlation (Figure 5A, blue), while most clusters showed no correlation (Figure 5A, black and Supplementary Figure 4). In support of this observation, whole-mount immunohistochemistry showed that OHC and IHC structures are normal in mice with good hearing (Figure 5E) compared to aged mice with poor hearing (Figure 5F), demonstrating that hair cell loss is a primary driver of hearing loss in CFW mice (Wu et al., 2020).

**Figure 5.**
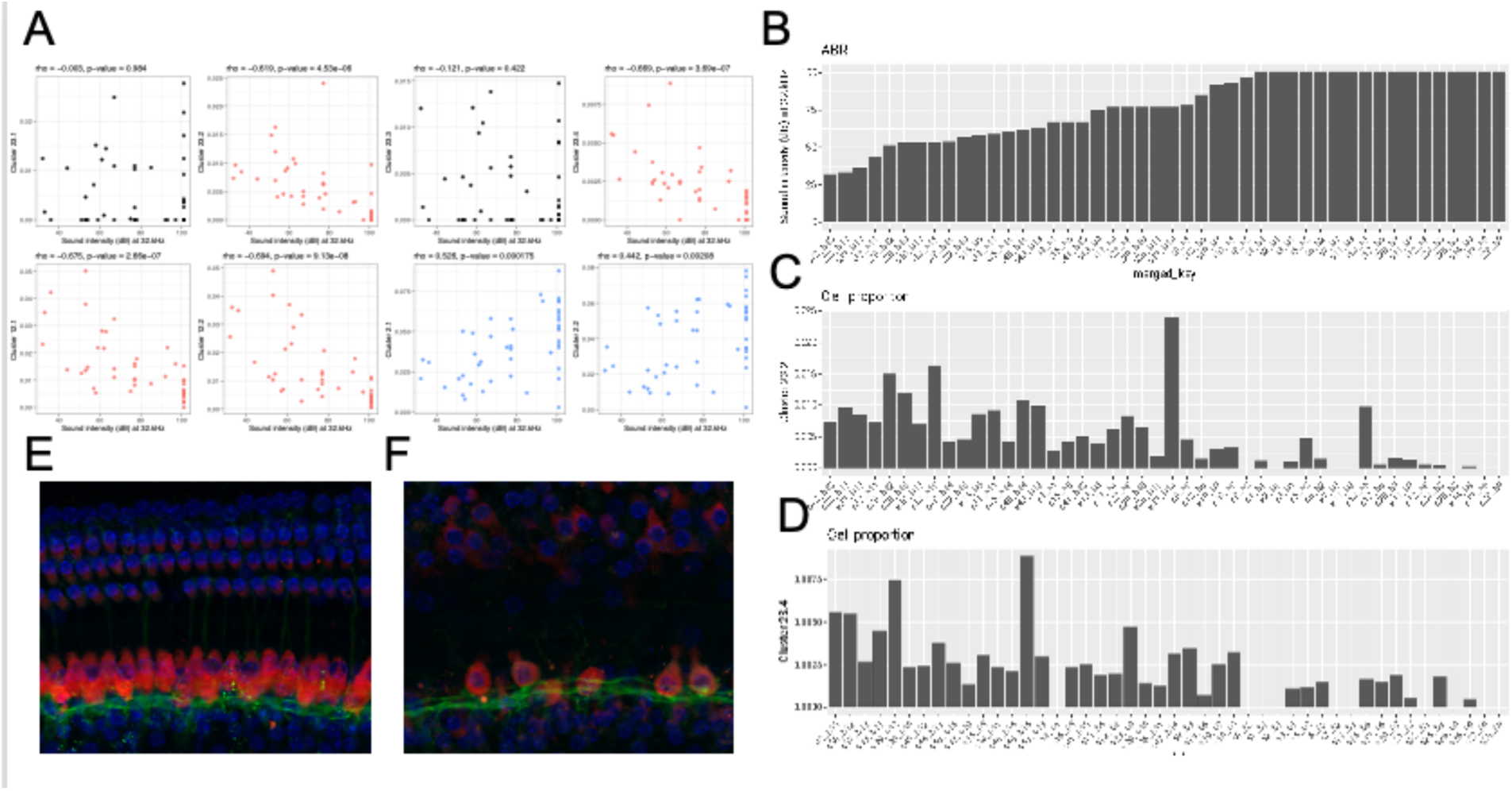
– Loss of specific cochlear cell types with age-related hearing loss. The proportion of each cell type in the 48 outbred mouse cochlea was compared against the ABR sound intensity score at different sound frequencies. A) Correlation between selected cell type proportions (Y-axis) and ABR sound intensity thresholds at 32 kHz (x-axis). Hair cell Clusters 23.2 and 23.4 and ganglion neuron Clusters 12.1 and 12.2 show significant inverse correlations with intensity threshold (red). Basal cell Clusters 2.1 and 2.2 show significant positive correlation with intensity threshold (blue). In contrast to the inverse correlation observed with Clusters 23.2 and 23.4, hair cell Clusters 23.1 and 23.3 show no correlation with intensity threshold. Spearman’s rho correlation and p-value are show at the top of each plot. B) Sound intensity threshold of individual mice ranked in order of hearing loss. C) Proportion of Cluster 23.2 cells in individual mice. D) Proportion of Cluster 23.4 cells in individual mice. Whole mount projections of organ of Corti corresponding to areas encoding 16 kHz frequency show that mice with all frequencies profound hearing loss (F) exhibit inner and outer hair cells loss that worsen through higher frequencies while the organ of Corti from mice with normal hearing (E) show intact structure and no apparent cells loss.

In comparing the cell type proportions with each other (Supplementary Figure 5), all cell types that show proportional loss with hearing loss are positively correlated with each other (e.g., Cluster 23.2 with Clusters 23.4, 12.1, and 12.2 (top row, left)), with the correlation strongest for cell types that are closely related to each other (e.g., Cluster 12.1 and 12.2, third row); all cell types that show proportional increases with hearing loss are positively correlated with each other (e.g., Cluster 2.1 with Clusters 2.2, 19.1, and 19.2 (fifth row, middle)); all of the cell types that show proportional increases with hearing loss are negatively correlated with cell types that show proportional loss with hearing (e.g., Cluster 2.1 with Clusters 23.2, 23.4, 12.1, and 12.2 (fifth row, left)); cell types that show no correlation with hearing loss show no correlation with cell types that show some correlation with hearing loss (e.g., Cluster 23.3 with Clusters 23.2, 23.4, 12.1, 12.2, 2.1, 2.2, 19.1, and 19.2 (bottom row, left and middle)). Interestingly, although the proportions of hair cell Clusters 23.1 and 23.3 do not correlate with hair cell clusters 23.2 and 23.4, they show strong correlation with each other. These results suggest that hearing loss is associated with specific concerted changes in cell populations in this outbred population and that two specific types of HC populations and two SGN populations are still preserved despite hearing loss.

## Discussion

In this manuscript we have described the first snRNA-seq study of the aged adult outbred mouse cochlea correlated with auditory physiological data. In contrast to a recent study, using scRNA-seq, which identified 27 cell types within the cochlea of a single mouse strain, C57BL/6J (Sun G et al.), we report the identification of 60 distinct cell types in this outbred mouse model using snRNA-seq to avoid the stress responses induced during the single cell processing procedure and to provide for a more unbiased representation of cell types from solid tissues (Bakken et al., 2018).

Presbycusis, ARHL, is the most common sensory disorder in man and is characterized by reduced hearing sensitivities and speech understanding, particularly in noisy environments. Classically described, there are four subtypes, sensory, neural, strial and mixed, each associated with a corresponding cellular senescence (Schuknecht and Gacek. 1993). The findings reported here confirm the complex nature of ARHL regarding the survival/senescence of specific cell types. It is well known that SGN numbers decrease with age and are responsible for mixed and sensory forms of hearing loss including their ultimate decline after synaptic ribbon loss (Fernandez et al., 2015). In this study, two of the four SGN clusters decreased with increasing hearing loss, and yet, two subtypes of SGN cells identified in this analysis did not. It is also well known that HC loss is a common finding in most acquired hearing loss, yet, of the four subtypes of HCs detected in our in-depth analysis, only two declined with hearing loss. We also observed that pillar and inner border cell populations declined with hearing loss resulting in a loss of cytoarchitecture in the organ of Corti. Elevated hearing thresholds were associated with loss of marginal cell clusters in the stria vascularis, the portion of the ear responsible for maintenance of the endocochlear potential. The increased number of basal cells and spindle cells positively correlating with higher hearing thresholds suggests a compensatory role for these cells in maintaining the endocochlear potential with aging (Hibino and Kurachi. 2006). The increased numbers of macrophages positively correlated with hearing loss likely represents an inflammatory response to the age-related cochlear stressors (Hough et al., 2022; Rai et al., 2020; Noble et al., 2022). Finally, our analysis identified seven separate clusters of fibrocytes with two out of the seven declining in numbers with hearing loss.

The use of snRNA-seq to identify and quantify cell types in the cochlea of elderly mice with varying levels of age-related hearing loss (ARHL) identified ten cell types whose proportions inversely correlated with hearing acuity, including two hair cell types (cochlear outer hair cell Slc26a5 and cochlear hair cell Ripor3), two spiral ganglion neuron types (cochlear spiral ganglion neuron C130073E24Rik Ntng1 and cochlear spiral ganglion neuron Mdga1 Tmem108), one Schwan cell (cochlear Schwan cell Pde1c Lama1), four border cell types (cochlear inner / outer sulcus Prss36 Hmcn1, cochlear border cell Dgkb Klhl14, cochlear pillar cell Mdm1, and cochlear phalageal cell Dgkb Nckap5), and one marginal cell type (cochlear marginal Stac Kcnq1) (Supplementary Table 3). Each of these distinct cell types is characterized by the expression of specific combinations of marker genes. Since these marker genes show very specific expression in the cell type that they mark, they may play a critical role in the function of that cell type. For example, the two marker genes for cochlear marginal Stac Kcnq1 cells are both involved in ion channel function that regulate plasma membrane potential (Wong King Yuen et al., 2017; Schroeder et al., 2000) and are likely to be essential for the proper functioning of the cell. Since these cell types are critical for maintaining hearing with age, it is possible that their marker genes would also be essential for hearing. Indeed three of the 19 marker genes for these ARHL-associated cell types show genetic associations with hearing deficits (Supplementary Table 3). Kvlqt1 (aka Kcnq1) -/-mice are completely deaf due to defects in inner ear development (Lee et al., 2000). Mutations in KVLQT1 in humans cause Jervell and Lange-Nielsen (JLN) syndrome, an inherited autosomal recessive disease characterized by a congenital bilateral deafness associated with QT prolongation and ventricular arrhythmias (Neyroud et al., 1997). Mutations in PDE1C are associated with an autosomal dominant form of nonsyndromic postlingual progressive deafness in human (Wang et al., 2018). Mutations in SLC26A5 (aka Prestin) are associated with familial nonsyndromic hearing loss in humans (Liu et al., 2003). Homozygous Slc26a5 mutant mice showed a loss of outer hair cell electromotility *in vitro* and a loss of cochlear sensitivity *in vivo* without disruption of mechanoelectrical transduction in outer hair cells (Liberman et al., 2002). The finding that some of these cell type specific marker genes show genetic association with hearing deficits supports the observation that loss of these specific cell types would lead to ARHL.

Interestingly, marker genes specific for other cell types that did not correlate with ARHL in our mouse model also show genetic associations with certain types of hearing deficits (Supplementary Table 4). For example, although the proportion of cochlear hair cell Ush2a C230072F16Rik did not diminish with ARHL, mutations in its marker gene USH2A causes Usher syndrome type IIa, an autosomal recessive disorder characterized by moderate to severe sensorineural hearing loss and retinitis pigmentosa (Eudy et al., 1998). These findings suggest that five other cochlear cell types identified in our study are involved in other types of hearing loss syndromes.

In conclusion, by combining information from manual marker gene annotation, gEAR reference data matching and MERFISH spatial transcriptomics analysis, 60 unique transcriptomic clusters were identified and annotated with cell type identities and specific marker gene characterization. Several of these specific cell types showed preferential loss in the aging cochlea that also correlated with quantitative measures of hearing loss. The genes specifically expressed in these cells could serve as candidate targets for novel therapeutics in the future.

## Materials and Methods

### Animals

Animals originated from the same stock of outbred mice Crl:CFW(SW)-US_P08 (CFW), maintained by Charles River Laboratories (CRL) in Portage, Michigan, were used in this study. All procedures were performed in accordance with guidelines from NIH and with approval by the Institutional Care and Use Committee (IACUC) at University of California San Diego (IACUC S17178).

### Auditory phenotyping/ Hearing Patterns determination

All CFW mice were subsequently aged until a final age of ten months. Within this period, phenotypic audiometric assessment using Auditory Brainstem Response (ABR) was completed at three time points: young adults (5-8 weeks), 6 months, and 10 months. Auditory phenotyping and determination of hearing patterns were previously detailed in (Du et al., 2022). Briefly, upon aging to the stated time points, anesthetized mice were presented to auditory signals as tone pips ranging from 20 dB to 100 dB SPL at the frequencies 4 kHz, 8 kHz, 12 kHz, 16 kHz, 24 kHz, and 32 kHz.

The detection distinctive ABR waveform at each frequency was used to determine the hearing thresholds and characterize ARHL within the CFW mice into one of eight distinct hearing patterns: Normal hearing; isolated mid-frequency (12 kHz or 16 kHz) loss; Moderate high-frequency (24 kHz or 32 kHz) loss; Severe high-frequency (only 32 kHz) loss; Severe high-frequency (24 kHz + 32 kHz) loss; Moderate all-frequency loss; Severe all-frequency loss; and Profound all-frequency loss.

### Isolation of cochlear tissue

A group of 48 mice were sacrificed and their inner ears were utilized to generate single nucleus RNA-seq transcriptomes. Mice were selected to represent the hearing pattern group described above based on the proportion of such pattern in the general CFW cohort. Special care was taken to complete the microdissection of the 2 cochleae from each mouse in less than 7 minutes while ensuring a minimum physical stretching on the organ of Corti. Samples were collected at the same time of day across individual and batches.

Following the ABR testing at 10 months age final time point, the anesthetized mice were decapitated, and their inner ears were quickly transferred into ice-cold DPBS buffer (Thermo Fisher Scientific) for microdissection. Tissue from utricule and saccule were carefully removed to avoid including unwanted vestibular cell types before removing the bony wall of the cochlea. The microdissected tissue from each mouse was pooled in a tube containing 1 ml of Iscove’s Modified Dulbecco’s Medium (IMDM) (Thermo Fisher Scientific).

### Single nucleus isolation and RNA sequencing

Immediately after isolation, bulk cochlear tissues in cell culture media were processed for snRNA-seq serially in sets of four specimens and kept on ice for the entire procedure. Dounce homogenization was performed to isolate individual nuclei from the bulk cochlear tissue followed by Fluorescent Automated Cell Sorting (FACS) to provide purified, intact nuclei for RNA-seq as detailed in (Krishnaswami 2016) with the following modifications. Briefly, a total of 5 mLs of Homogenization Buffer with Propidium Iodide (PI) (1.5 µM, Thermo Fisher cat. #P3566) and Calcien, AM (1.0 µM) (Thermo Fisher cat. #C1430) was used to stain intact nuclei and exclude whole cells from being sorted. The cochlear tissue in cell culture media was briefly centrifuged to remove the supernatant to approximately 100 µL volume and the remaining tissue transferred using a 1 mL wide bore pipette tip into 1 mL of chilled Homogenization Buffer in a Dounce Homogenizer (Wheaton cat. #357538) stored on ice. A total of 10 strokes with the loose piston followed by 14 strokes with the tight piston was followed by filtration through a series of two cell strainers (Becton Dickenson Falcon cat. #352235) and loading onto a Beckton Dickenson FACS Aria II cell sorter with a 70 mm nozzle. A chilled 96-well plate was used with 4 µL of Homogenization Buffer placed in wells A1 to A4 as a destination for sorted nuclei. FACS gating of nuclei were accomplished with forward and side scatter doublet nuclei gating exclusion, green channel (Calcien, AM) exclusion, and triggered on the red (PI) channel with approximately 1-7% of the total event population consisting of intact nuclei. A total of 36,000 sorted nuclei from each of the 4 samples were sorted into wells A1 to A4 with a final volume of approximately 60 µL after completion of sorting.

Isolated nuclei in the 4 wells were kept on ice and processed for RNA-seq using the Chromium Next GEM Chip G Single Cell Kit (10X Genomics cat. #1000120) according to manufactures instructions with the following modifications. A total of 43.2 µL of sorted volume containing approximately 26,000 nuclei was added to 31.8 µL of the Single Cell Master mix for a total of 75 µL, followed by loading of 70 µL (approx. 24,267 nuclei) onto the Next GEM chip. cDNAs were amplified for 13 cycles. Recovered cDNAs were quality controlled by loading 1 µL of cDNA onto an Agilent Bioanalyzer using a High Sensitivity DNA chip kit (cat. #5067-4626), followed by library prep according to manufacturer’s instructions for dual index barcoded libraries.

The nuclei isolation, sorting, and Next Gem chip cDNA synthesis procedures were repeated 12 more times for a total of 48 cDNA library samples. A total of 48 single nuclei barcoded RNA-seq libraries were subsequently generated from the cDNAs using the Single Index Kit T Set A (10X Genomics cat. #1000213) and 16 cycles of library amplification followed by purification according to manufactures instructions. A 1 µL loading of a 10-fold dilution of each library was performed for quality control and quantification on an Agilent High Sensitivity chip. Equimolar amounts (6 nM) of a subset of 16 libraries were pooled and diluted to create a final loading of a 300 pM library pool onto a dual-lane NovaSeq 6000 (Illumina, Inc.) using the S2 Reagent Kit v1.5 Paired End 2×50 base (100 cycle) sequencing kit (Illumina cat. #20028316). Sequencing parameters were set for R1 at 28 cycles, I7 Index at 8 cycles, I5 index at 0 cycles, and Read 2 at 91 cycles for a calculated R2 read count of 25,600 per cell. Two more NovaSeq 6000 S2 runs of 16 library pools were subsequently loaded for a total of 3 runs to complete the RNA-seq of all 48 samples. A range of X to Y reads per cell were generated.

### Read alignment

The cellranger count command with the –include-intron option from the 10x Cell Ranger 5.0.1 package (Zheng 2017) was used to align reads and count barcodes and UMIs. Reads were aligned to the Cell Ranger reference package refdata-gex-mm10-2020-A.

### Quality control

Data quality was assessed using the number of nuclei per sample, the number of UMIs (library size), the number of genes detected and the number of mitochondrial genes per nuclei. Nuclei were removed if they had fewer than 1000 UMIs and/or fewer than 500 detected genes. Nuclei that had the number of detected mitochondrial genes tagged as outliers by the isOutlier function from the scuttle package were also removed (McCarthy 2017).

### Normalization, feature selection and dimensionality reduction

UMI counts were normalized using logNormCounts from the scuttle package. Highly variable genes were selected using modelGeneVar and getTopHVGs functions from the scran package (Lun 2016). PCA was performed using the runPCA function from the scater package. Nineteen principal components were used for clustering, which accounted for 50% of the variance.

### Clustering and doublet detection

To minimize the inclusion of droplets that have more than one nucleus, putative doublets were identified using the scDblFinder package (Germain 2021). Since scDblFinder uses cluster information to predict doublets, nuclei were first clustered using the Leiden community detection algorithm as implemented in the leidenalg python package (Traag 2018). First, a shared nearest-neighbor graph was constructed with buildSNNGraph from the scran package (McCarthy 2017). Then the leidenalg.find_partition function with partition_type set to ModularityVertexPartition was called. Finally, sclDblFinder was called using the resultant cluster labels. About 6% of the droplets were predicted to be doublets and removed from downstream analysis (Supplementary Figure 6). The remaining nuclei were re-processes repeating quality control, normalization, feature selection, dimensionality reduction and clustering.

### Visualization

The clustered data set was visualized by running runUMAP and runTSNE from the scater package (McCarthy 2017). For runUMAP, n_neighbors was set to 100; for runTSNE, perplexity was set to 100.

### Sub-clustering

The initial clustering using Leiden community detection yielded 26 clusters. To determine if any of these clusters should undergo another round of clustering, two methods were used: the silhouette width, which compares the average distance of a nucleus to all other nuclei within the same cluster to the average distance to nuclei in the nearest neighboring cluster^70^ [Amezquita 2021 section 5.2.2], and manual inspection. Supplementary Figure 7 shows an example of this analysis where Clusters 1, 10, and 17 contain separate areas of nuclei with negative silhouette widths colored in red as candidates for sub-clustering. Other candidate clusters were detected by manual inspection of the tSNE embeddings (Figure 2A), for example, Clusters 20 and 23, which appear to have two distinct “islands” suggesting the need for sub-clustering. Supplementary Figure 8 shows examples of the resulting subclusters. By applying those criteria on the initial clusters, 13 were retained and 13 went through a second round of sub-clustering, which brought the total number of transcriptomic clusters identified to 60. The final clustered cell-by-gene expression matrix can be found at the gEAR resource -https://umgear.org/p?l=c2acc279.

### Cluster markers

NS-Forest was used to identify the minimum number of necessary and sufficient markers for each cluster (Aevermann 2021). NS-Forest was performed with the number of trees set to 1000 and the number of genes to test set to 6.

### Multiplexed error-robust fluorescence in situ hybridization (MERFISH) on the cochlear tissue

MERFISH samples were prepared in accordance with company instructions (Vizgen, Cambridge, MA, USA). Briefly, C57Bl/6J mice cochleae of postnatal day 5 and 2 months were harvested and fixed with 4% paraformaldehyde (PFA) in 0.1 M phosphate-buffered saline (PBS; pH 7.4) at 4 °C overnight. Samples were dehydrated in graded-sucrose series (10% and 20% for 30 min, and 30% for overnight at 4 °C) with RNase inhibitor [New England Biolabs (NEB), M0314L, Ipswich, MA, USA]. Samples were placed in a cryomold and embedded with O.C.T. compound (Sakura Finetek, Torrance, CA, USA), then frozen with dry ice/ethanol bath. The embedded tissue was sectioned into 10μm thick slices using a Leica CM1860 (Leica Biosystems, Nussloch, Germany) cryostat, and 3 to 5 cochlear mid-modiolar sections were mounted onto a center of MERSCOPE slide glass (Vizgen, #20400001, USA). The mounted sections were washed with 0.1 M PBS, then permeabilized in 70% ethanol at 4 °C for 24 h. The cell boundary staining was performed by using a primary antibody mix (Vizgen, #20300010, USA), followed by a secondary antibody mix (Vizgen, #20300011, USA) for 1 h at 23 ℃, respectively. Stained samples were incubated with an encoding probe [MERSCOPE 140 Gene Panel Mix (Vizgen, #20300006, USA)] for 36 hours at humidified 37 ℃ cell culture incubator. After post-encoding hybridization wash with formamide wash buffer (Vizgen, #20300002, USA), samples were embedded with a gel embedding solution [gel embedding premix (Vizgen, #20300004, USA), 5 ml; 10% ammonium persulfate solution (Millipore-Sigma, 09913-100G, Burlington, MA, USA), 25 µl; N,N,N’,N’-tetramethylethylenediamine (Millipore-Sigma, T7024-25ML, USA), 2.5 µl]. For tissue clearing, samples were incubated in digestion premix (Vizgen, #20300005, USA) with RNase inhibitor (NEB, USA) for 1 h at 23 °C, followed by clearing premix [clearing premix (Vizgen, #20300003, USA), 5ml; proteinase K (NEB, P8107S, USA), 501l] for 48 h at humidified 37 ℃ cell culture incubator. After the tissue became transparent, samples were washed with the wash buffer (Vizgen, #20300001, USA) and incubated with 4′,6-diamidino-2-phenylindole (DAPI) and polythymine (polyT) staining reagent (Vizgen, #20300021, USA) for 15 min with agitation. Images were taken by using MERSCOPE (Vizgen, USA). DAPI / polyT and cell-boundary staining 2 was utilized for the cell segmentation parameter respectively, then image processing analysis was done on the MERSCOPE. The images were visualized and analyzed on the MERSCOPE Visualizer (Vizgen, USA).

## ACKNOWLEDGEMENTS

Research reported in this publication was supported by NIDCD of the National Institutes of Health under award number: R01DC018566.

---

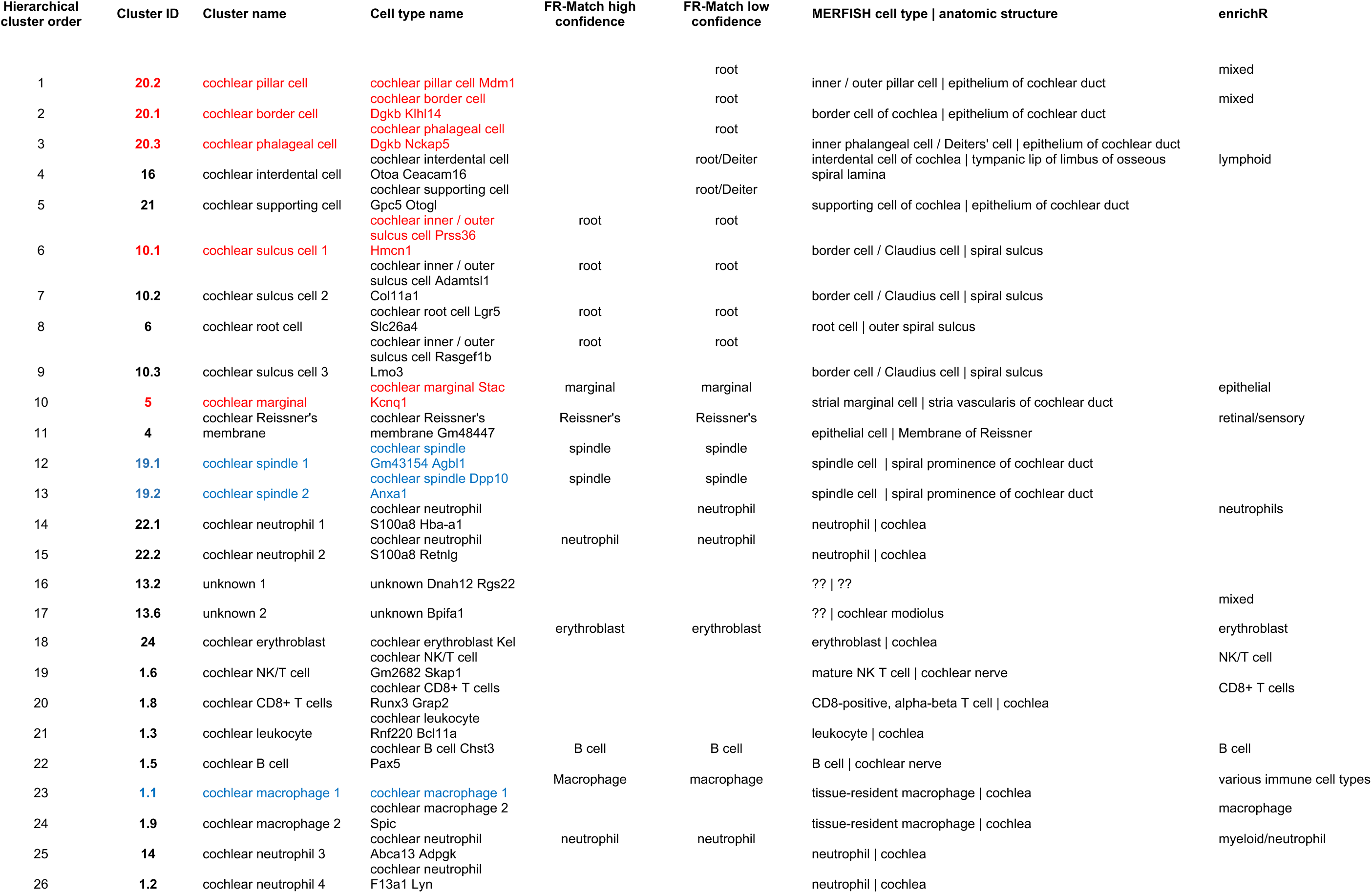
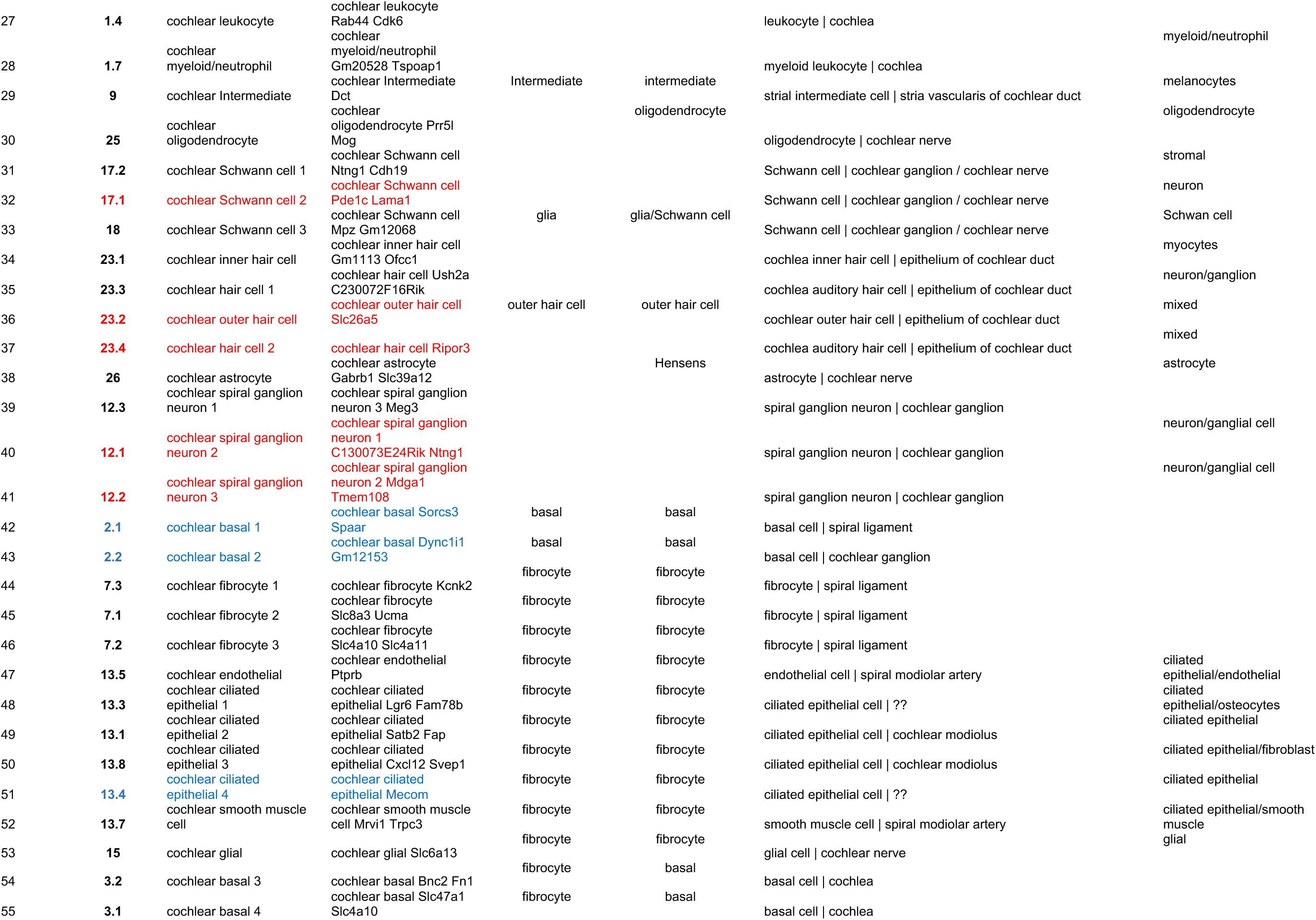
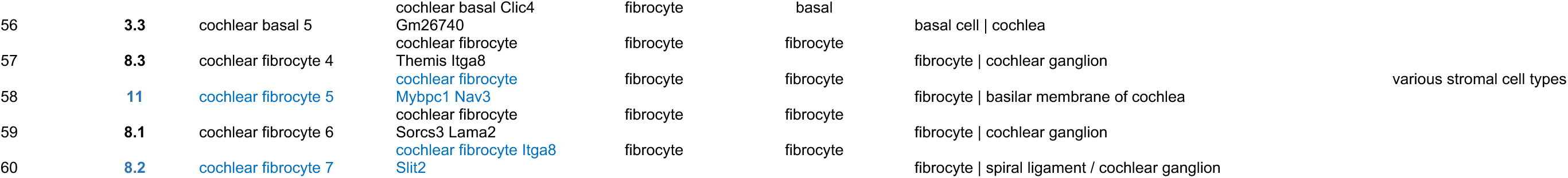

## Notes

### Competing Interest Statement

The authors have declared no competing interest.

### Summary of Updates

The Manuscript Title, Figure 4 revised, author list updated; Supplemental files updated

