## Supplementary Figures for "Cochlear transcriptome analysis of an outbred mouse population (CFW)"

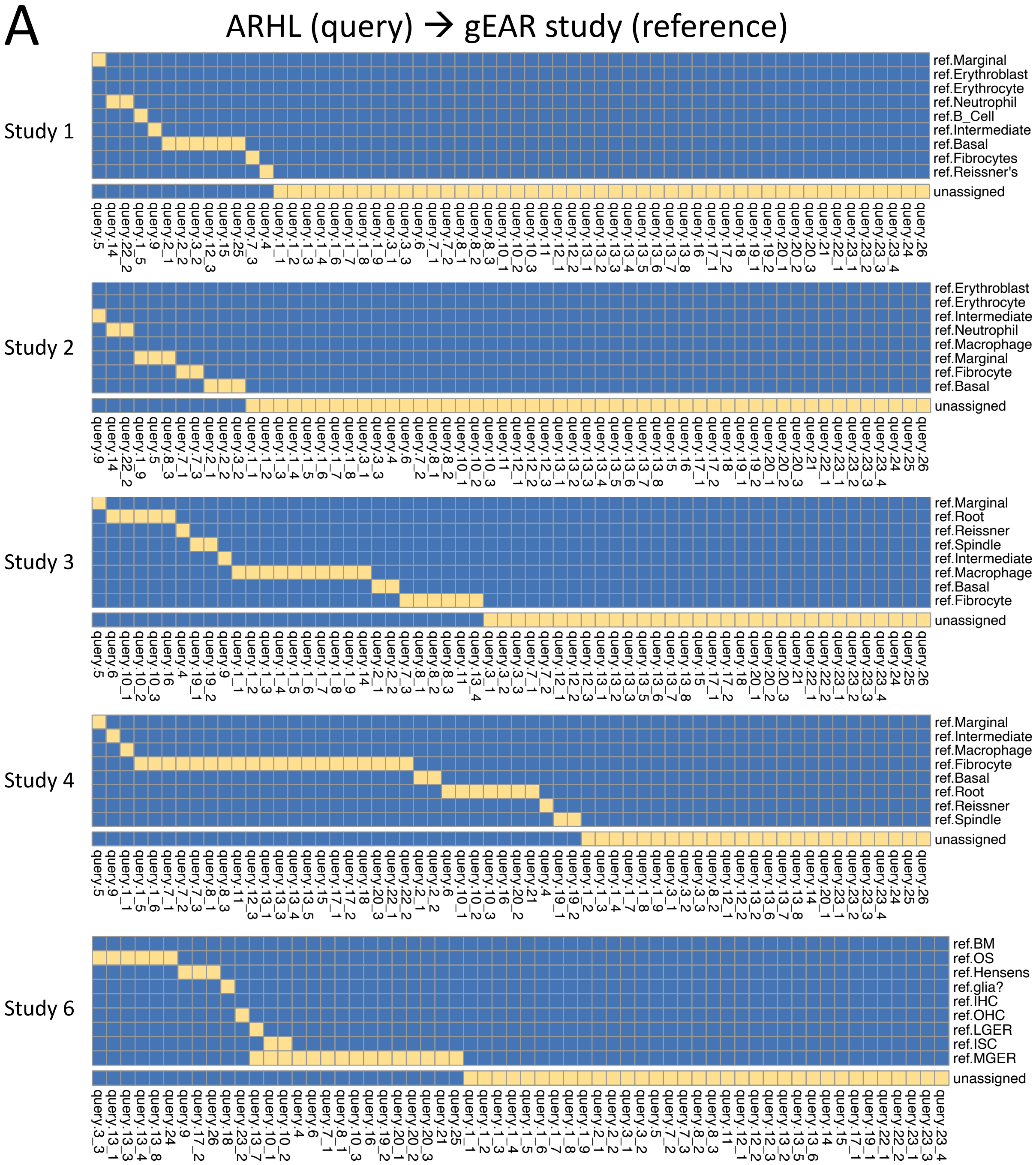

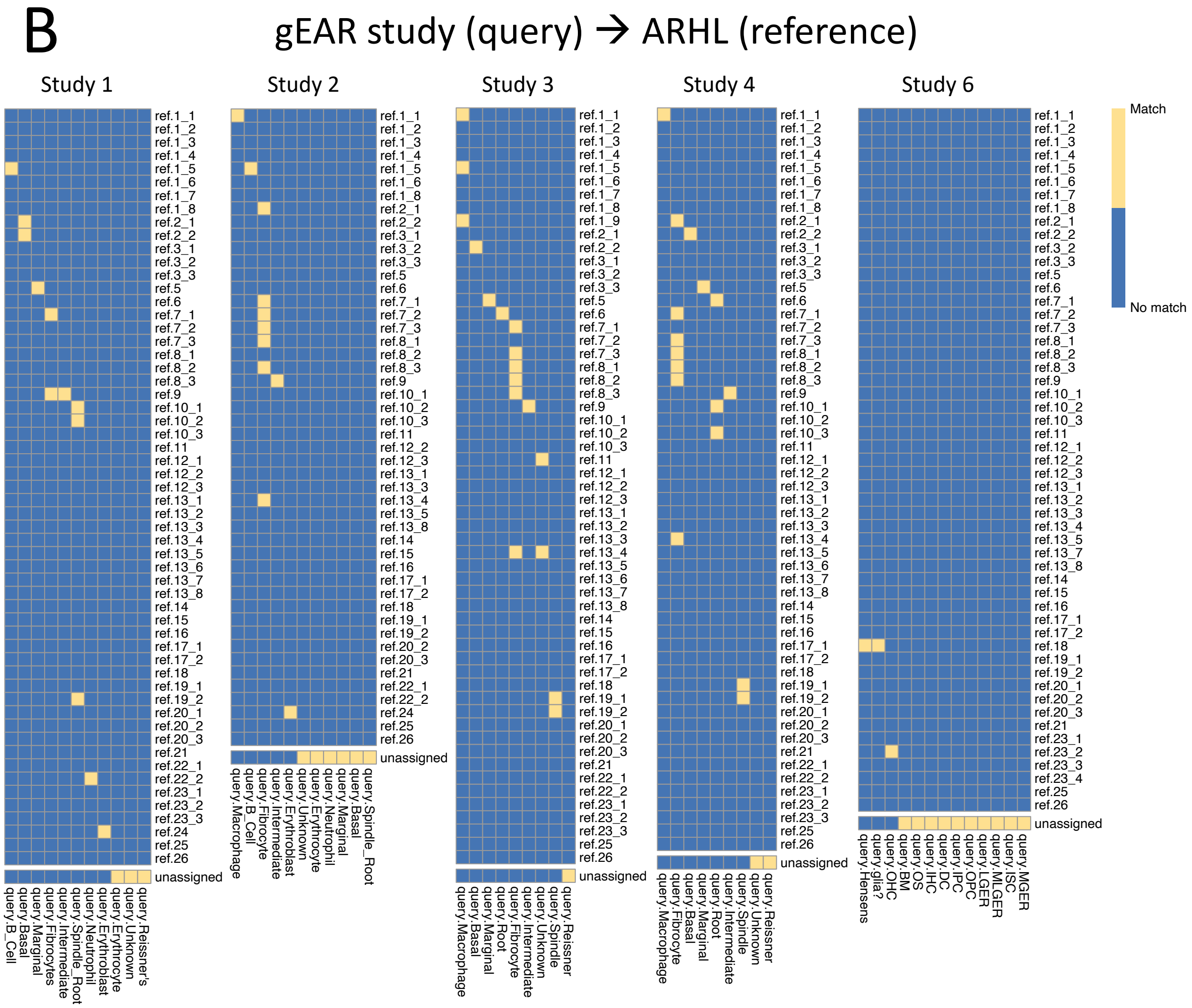


**Supplementary Figure 1 – Matching ARHL cell types with gEAR study cell types.** sc/snRNA-seq data from five relevant studies (Supplementary Table 2) posted on the gEAR resource website (https://umgear.org) were used to determine if the same cell types have been identified in the ARHL snRNA-seq study reported here. Matching was performed using the cluster-to-cluster matching version of FR-Match [Zhang 2022]; positive matches are indicated by a yellow box. A) NS-Forest [Aevermann 2021] was used to select marker genes for each of the cell types reported in the five gEAR studies used as the reference datasets for matching by the ARHL dataset as the query. B) ARHL NS-Forest marker genes (Figure 3 and Supplementary Table 1) were used for matching the gEAR study query data to the ARHL reference.

Supplemental Fig.2

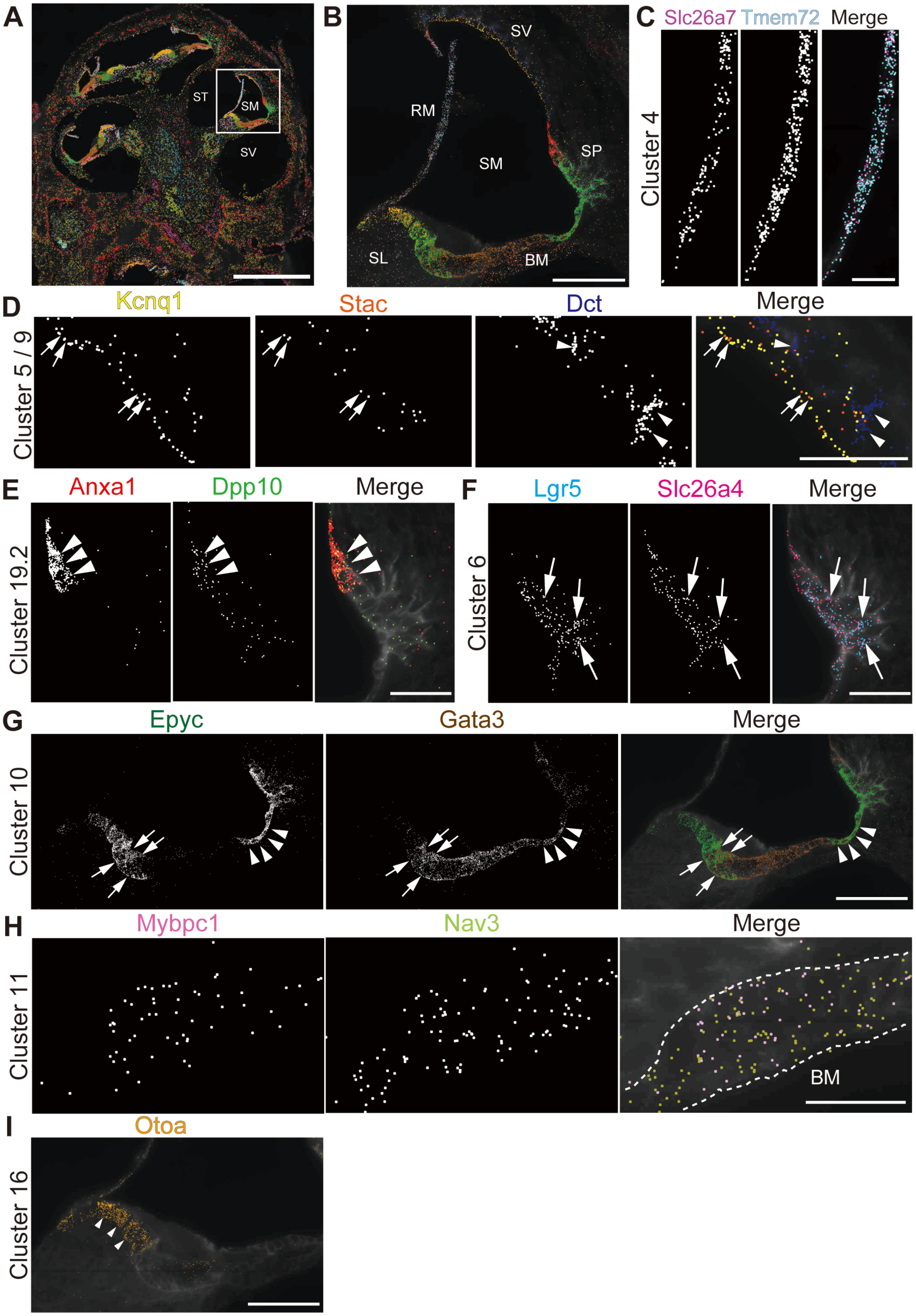

**Supplementary Figure 2 – Spatial mapping marker-gene expression across cochlear section by MERFISH.** Distribution of marker genes on the cochlear section at P4 using MERFISH. **A)** MERFISH image of the cochlear mid-modiolar cryosection with selected marker genes for clusters involved in the scala media: *Slc26a7* (purple) and *Tmem72* (light blue) for Cluster4 of Reisner membrane; *Kcnq1* (yellow) and *Stac* (orange) for Cluster 5 of marginal cell; *Lgr5* (blue) and *Slc26a4* (magenta) for Cluster 6 of root cell; *Dct* (dark blue) for Cluster 9 of intermediate cell; *Epyc* (dark green) and *Gata3* (brown) for Cluster 10 of sulcus cell; Otoa (kahaki) for Cluster 16 of interdental cell; *Mybpc1* (pink) and *Nav3* (light green) for Cluster 11 of fibrocyte; *Anxa1*(red) and *Dpp10*(green) for Cluster 19.2 of spindle cell. Note that the same color pattern was observed among different cochlea turs. ST; scala tympani, SM; scala media, SV; scala vestibuli. **B)** A high magnification view of the white boxed region in (A). BM; basilar membrane, RM; Reisner membrane, SL; spiral limbus, SM; scala media, SP; spiral prominence, SV; stria vascularis. **C)** A high magnification view of Reisner membrane region. **D)** Marginal cell (Cluster5, arrow) and intermediate cell (Cluster9, arrowhead) clusters in the stria vascularis. **E & F)** Spindle cell (Cluster 19.2, arrow heads) and root cell (Cluster 5, arrows) clusters in the spiral prominence. **G)** Cluster10 of inner (arrow) and outer sulcus (arrowhead) cell in the organ of Corti. **H)** Cluster 11 of fibrocyte in the connective tissue layer of the basilar membrane depicted by white dots. **I)** Cluster 16 of interdental cells in the spiral limbus. **Scale bar:** 500μm (A), 100 μm (B, G, I), 25μm (C-F, H)

Supplemental Fig.3

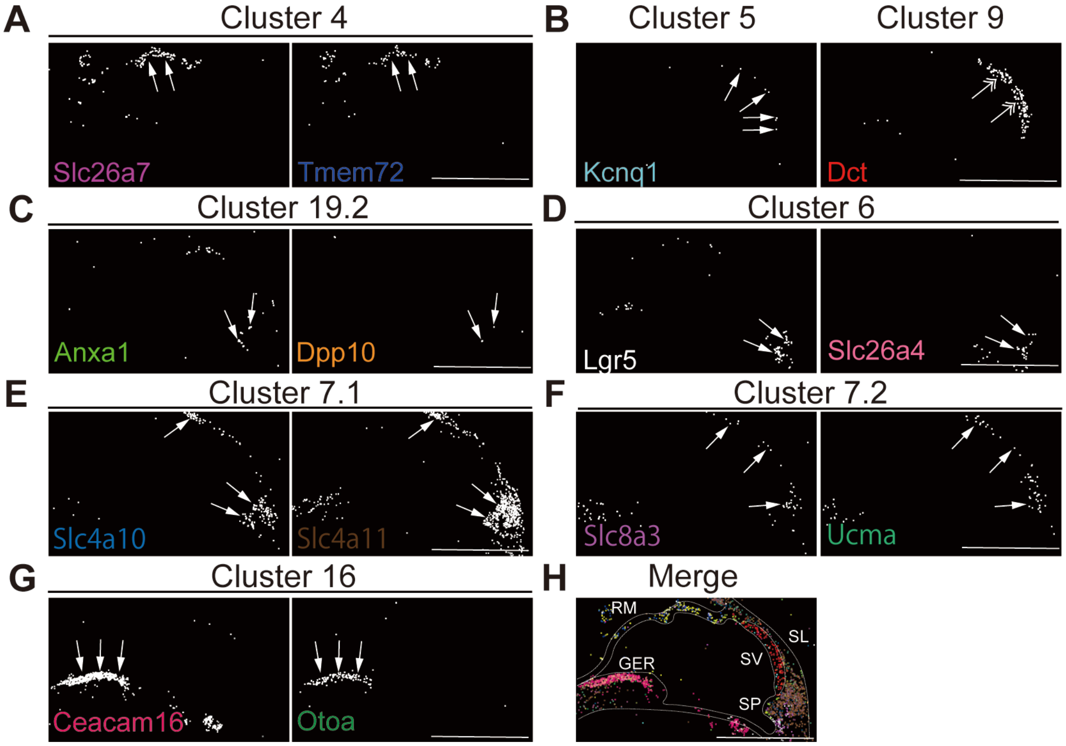


**Supplementary Figure 3 – Spatial mapping marker-gene expression across cochlear section by MERFISH in adult cochlea**

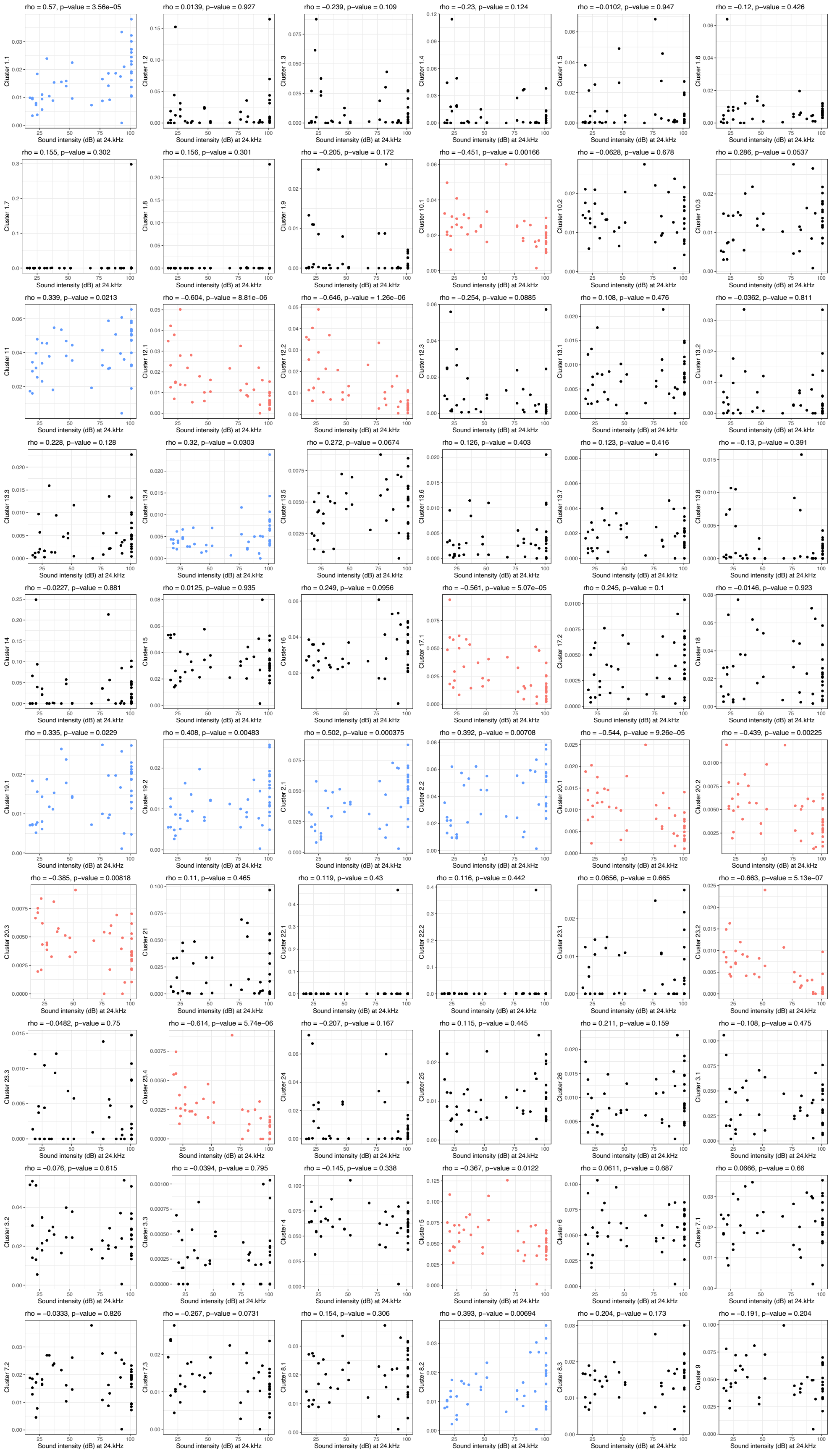
rho = 0.57, p−value = 3.56e−05
Cluster 1.1
Sound intensity (dB) at 24.kHz
rho = 0.0139, p−value = 0.927
Cluster 1.2
Sound intensity (dB) at 24.kHz
rho = −0.239, p−value = 0.109
Cluster 1.3
Sound intensity (dB) at 24.kHz
rho = −0.23, p−value = 0.124
Cluster 1.4
Sound intensity (dB) at 24.kHz
rho = −0.0102, p−value = 0.947
Cluster 1.5
Sound intensity (dB) at 24.kHz
rho = −0.12, p−value = 0.426
Cluster 1.6
Sound intensity (dB) at 24.kHz
rho = 0.155, p−value = 0.302
Cluster 1.7
Sound intensity (dB) at 24.kHz
rho = 0.156, p−value = 0.301
Cluster 1.8
Sound intensity (dB) at 24.kHz
rho = −0.205, p−value = 0.172
Cluster 1.9
Sound intensity (dB) at 24.kHz
rho = −0.451, p−value = 0.00166
Cluster 10.1
Sound intensity (dB) at 24.kHz
rho = −0.0628, p−value = 0.678
Cluster 10.2
Sound intensity (dB) at 24.kHz
rho = 0.286, p−value = 0.0537
Cluster 10.3
Sound intensity (dB) at 24.kHz
rho = 0.339, p−value = 0.0213
Cluster 11
Sound intensity (dB) at 24.kHz
rho = −0.604, p−value = 8.81e−06
Cluster 12.1
Sound intensity (dB) at 24.kHz
rho = −0.646, p−value = 1.26e−06
Cluster 12.2
Sound intensity (dB) at 24.kHz
rho = −0.254, p−value = 0.0885
Cluster 12.3
Sound intensity (dB) at 24.kHz
rho = 0.108, p−value = 0.476
Cluster 13.1
Sound intensity (dB) at 24.kHz
rho = −0.0362, p−value = 0.811
Cluster 13.2
Sound intensity (dB) at 24.kHz
rho = 0.228, p−value = 0.128
Cluster 13.3
Sound intensity (dB) at 24.kHz
rho = 0.32, p−value = 0.0303
Cluster 13.4
Sound intensity (dB) at 24.kHz
rho = 0.272, p−value = 0.0674
Cluster 13.5
Sound intensity (dB) at 24.kHz
rho = 0.126, p−value = 0.403
Cluster 13.6
Sound intensity (dB) at 24.kHz
rho = 0.123, p−value = 0.416
Cluster 13.7
Sound intensity (dB) at 24.kHz
rho = −0.13, p−value = 0.391
Cluster 13.8
Sound intensity (dB) at 24.kHz
rho = −0.0227, p−value = 0.881
Cluster 14
Sound intensity (dB) at 24.kHz
rho = 0.0125, p−value = 0.935
Cluster 15
Sound intensity (dB) at 24.kHz
rho = 0.249, p−value = 0.0956
Cluster 16
Sound intensity (dB) at 24.kHz
rho = −0.561, p−value = 5.07e−05
Cluster 17.1
Sound intensity (dB) at 24.kHz
rho = 0.245, p−value = 0.1
Cluster 17.2
Sound intensity (dB) at 24.kHz
rho = −0.0146, p−value = 0.923
Cluster 18
Sound intensity (dB) at 24.kHz
rho = 0.335, p−value = 0.0229
Cluster 19.1
Sound intensity (dB) at 24.kHz
rho = 0.408, p−value = 0.00483
Cluster 19.2
Sound intensity (dB) at 24.kHz
rho = 0.502, p−value = 0.000375
Cluster 2.1
Sound intensity (dB) at 24.kHz
rho = 0.392, p−value = 0.00708
Cluster 2.2
Sound intensity (dB) at 24.kHz
rho = −0.544, p−value = 9.26e−05
Cluster 20.1
Sound intensity (dB) at 24.kHz
rho = −0.439, p−value = 0.00225
Cluster 20.2
Sound intensity (dB) at 24.kHz
rho = −0.385, p−value = 0.00818
Cluster 20.3
Sound intensity (dB) at 24.kHz
rho = 0.11, p−value = 0.465
Cluster 21
Sound intensity (dB) at 24.kHz
rho = 0.119, p−value = 0.43
Cluster 22.1
Sound intensity (dB) at 24.kHz
rho = 0.116, p−value = 0.442
Cluster 22.2
Sound intensity (dB) at 24.kHz
rho = 0.0656, p−value = 0.665
Cluster 23.1
Sound intensity (dB) at 24.kHz
rho = −0.663, p−value = 5.13e−07
Cluster 23.2
Sound intensity (dB) at 24.kHz
rho = −0.0482, p−value = 0.75
Cluster 23.3
Sound intensity (dB) at 24.kHz
rho = −0.614, p−value = 5.74e−06
Cluster 23.4
Sound intensity (dB) at 24.kHz
rho = −0.207, p−value = 0.167
Cluster 24
Sound intensity (dB) at 24.kHz
rho = 0.115, p−value = 0.445
Cluster 25
Sound intensity (dB) at 24.kHz
rho = 0.211, p−value = 0.159
Cluster 26
Sound intensity (dB) at 24.kHz
rho = −0.108, p−value = 0.475
Cluster 3.1
Sound intensity (dB) at 24.kHz
rho = −0.076, p−value = 0.615
Cluster 3.2
Sound intensity (dB) at 24.kHz
rho = −0.0394, p−value = 0.795
Cluster 3.3
Sound intensity (dB) at 24.kHz
rho = −0.145, p−value = 0.338
Cluster 4
Sound intensity (dB) at 24.kHz
rho = −0.367, p−value = 0.0122
Cluster 5
Sound intensity (dB) at 24.kHz
rho = 0.0611, p−value = 0.687
Cluster 6
Sound intensity (dB) at 24.kHz
rho = 0.0666, p−value = 0.66
Cluster 7.1
Sound intensity (dB) at 24.kHz
rho = −0.0333, p−value = 0.826
Cluster 7.2
Sound intensity (dB) at 24.kHz
rho = −0.267, p−value = 0.0731
Cluster 7.3
Sound intensity (dB) at 24.kHz
rho = 0.154, p−value = 0.306
Cluster 8.1
Sound intensity (dB) at 24.kHz
rho = 0.393, p−value = 0.00694
Cluster 8.2
Sound intensity (dB) at 24.kHz
rho = 0.204, p−value = 0.173
Cluster 8.3
Sound intensity (dB) at 24.kHz
rho = −0.191, p−value = 0.204
Cluster 9
Sound intensity (dB) at 24.kHz

**Supplementary Figure 4 - Loss of specific cochlear cell types with age-related hearing loss.** The proportion of each cell type in the 48 outbred mouse cochlea was compared against the ABR sound intensity score at different sound frequencies. Correlations between cell type cluster proportions (Y-axis) and ABR sound intensity thresholds (dB) at 24 kHz (x-axis) are shown. Clusters showing a significant inverse correlation between cell type proportions and sound intensity threshold are colored red; clusters showing a significant positive correlation with intensity threshold are colored blue; clusters with no significant correlation with intensity threshold are colored black. Spearman's rho correlations and p-values are show at the top of each plot.

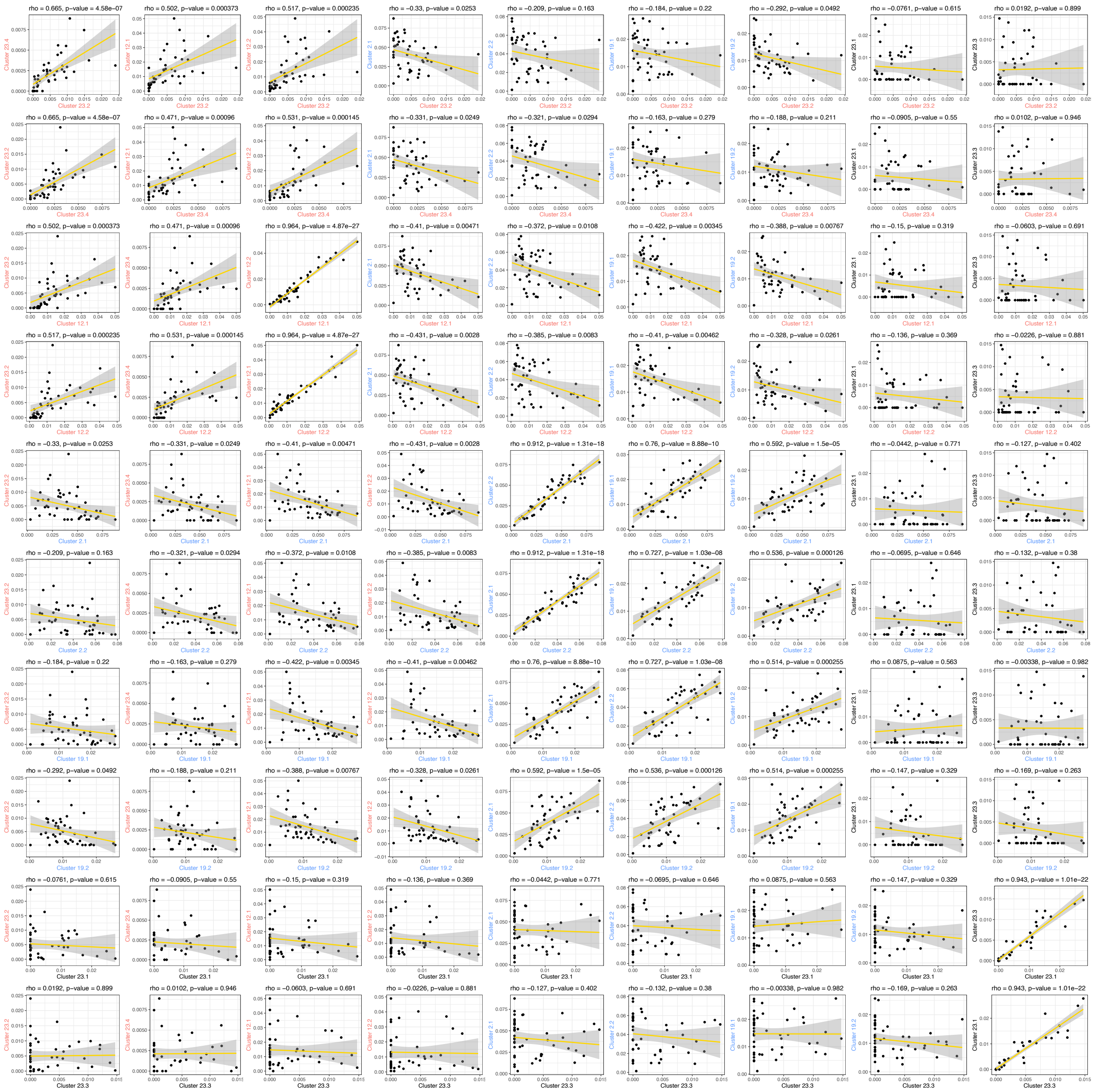


**Supplementary Figure 5 – Correlations between cell proportions.** The correlations between cell cluster proportions for selected pairs of clusters is shown. Clusters that showed a significant inverse correlation with intensity threshold in Supplementary Figure 3 are colored red; clusters with a significant positive correlation with intensity threshold in Supplementary Figure 3 are colored blue. Clusters with no significant correlation with intensity threshold in Supplementary Figure 3 are colored black. Spearman's rho correlations and p-values are show at the top of each plot.

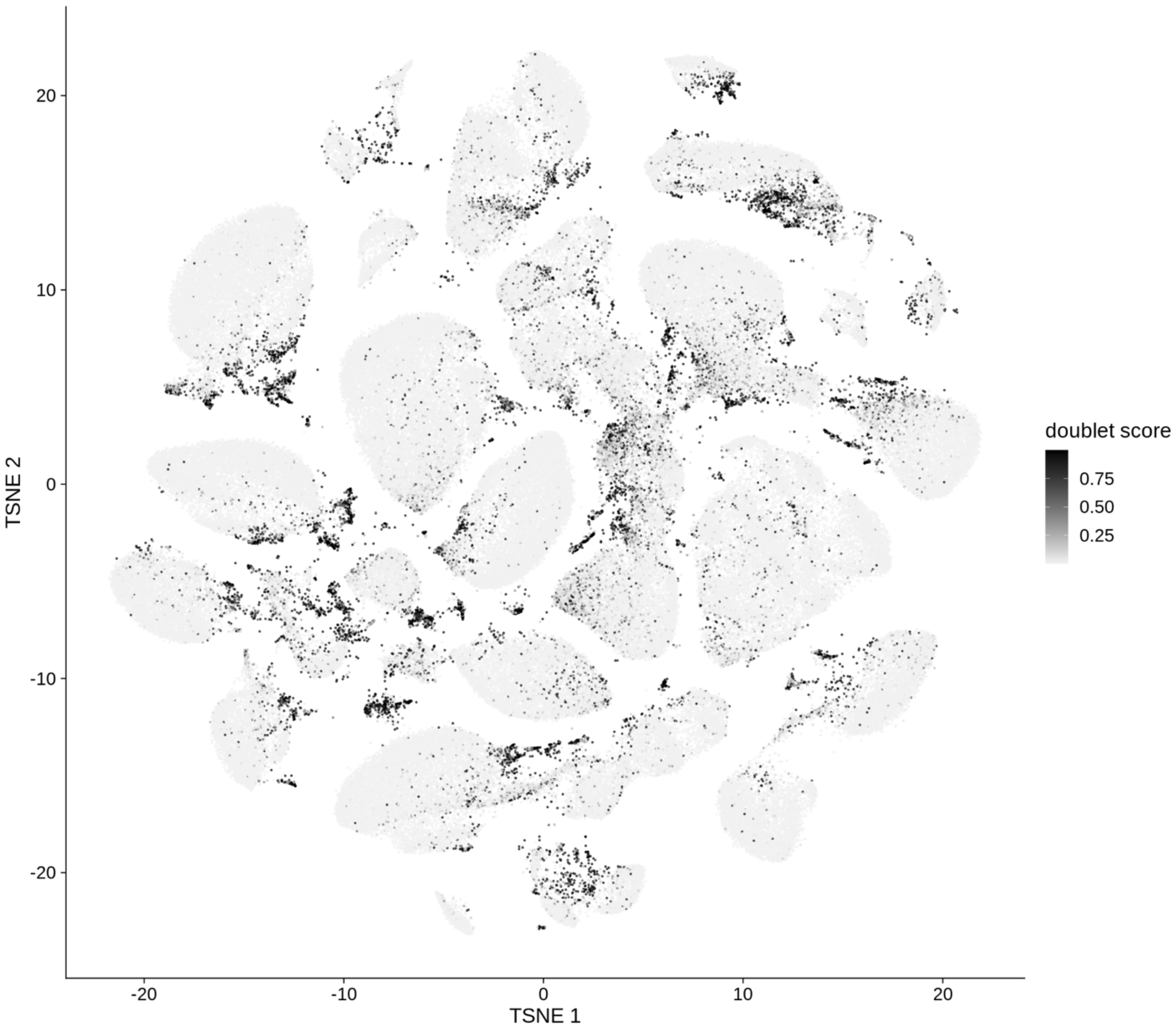


**Supplementary Figure 6 – snRNA-seq single nucleus tSNE embedding of putative doublets.** Nuclei were first clustered using the Leiden community detection algorithm as implemented in the leidenalg python package [Traag 2018]. First, a shared nearest-neighbor graph was constructed with buildSNNGraph from the scran package [McCarthy 2017]. Then the leidenalg.find_partition function with partition_type set to ModularityVertexPartition was called. Finally, sclDblFinder was called using the resultant cluster labels. Nuclei are colored by the resulting doublet score. About 6% of the nuclei droplets were predicted to be doublets and removed from downstream analysis.

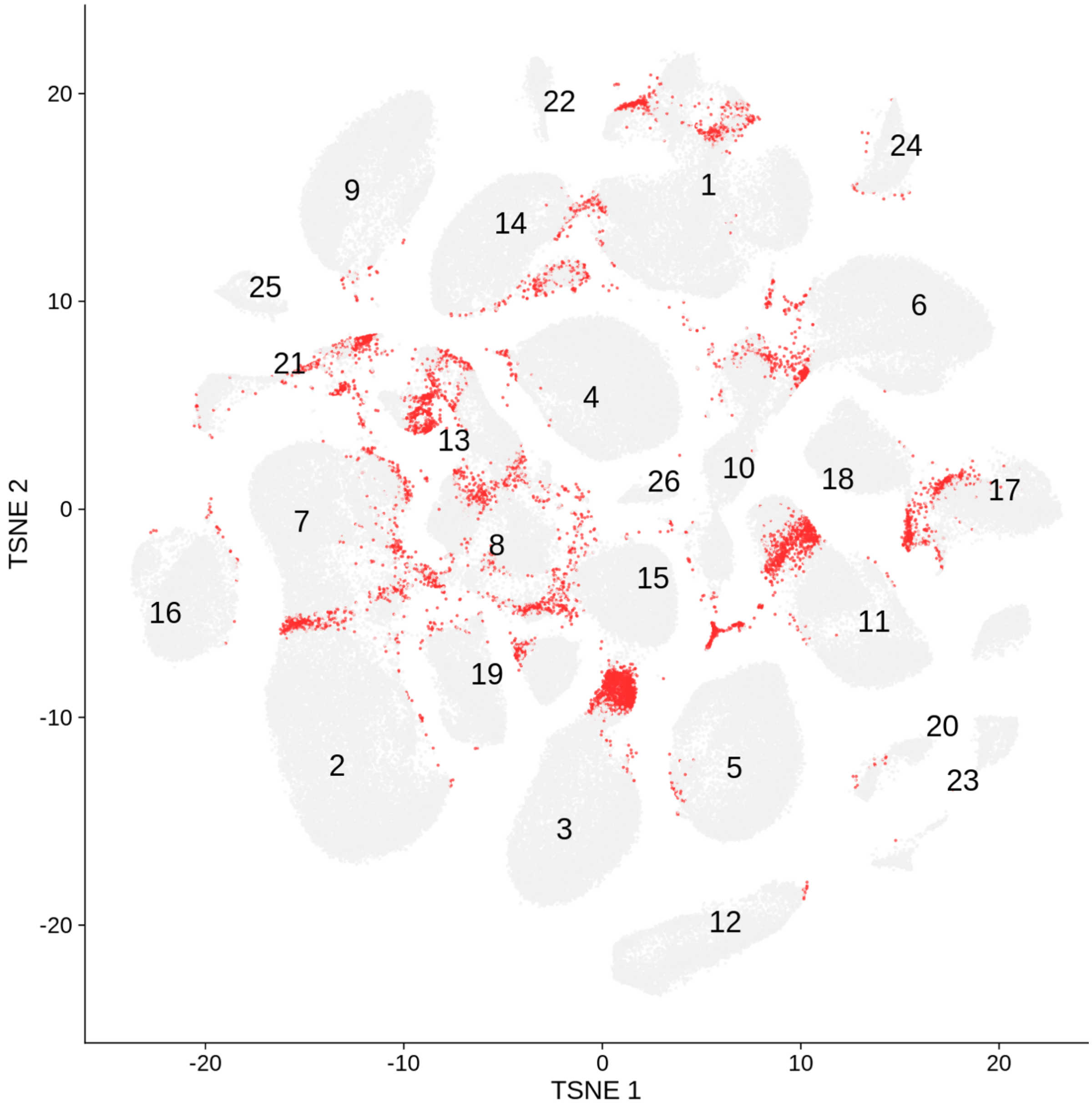


**Supplementary Figure 7 – Silhouette analysis of initial clustering results.** tSNE embedding of the initial set of 26 clusters identified by Leiden community detection is shown. Nuclei with negative silhouette widths are colored red. Silhouette width compares the average distance of a nucleus to all other nuclei within the same cluster to the average distance to nuclei in other clusters [Amezquita 2021].

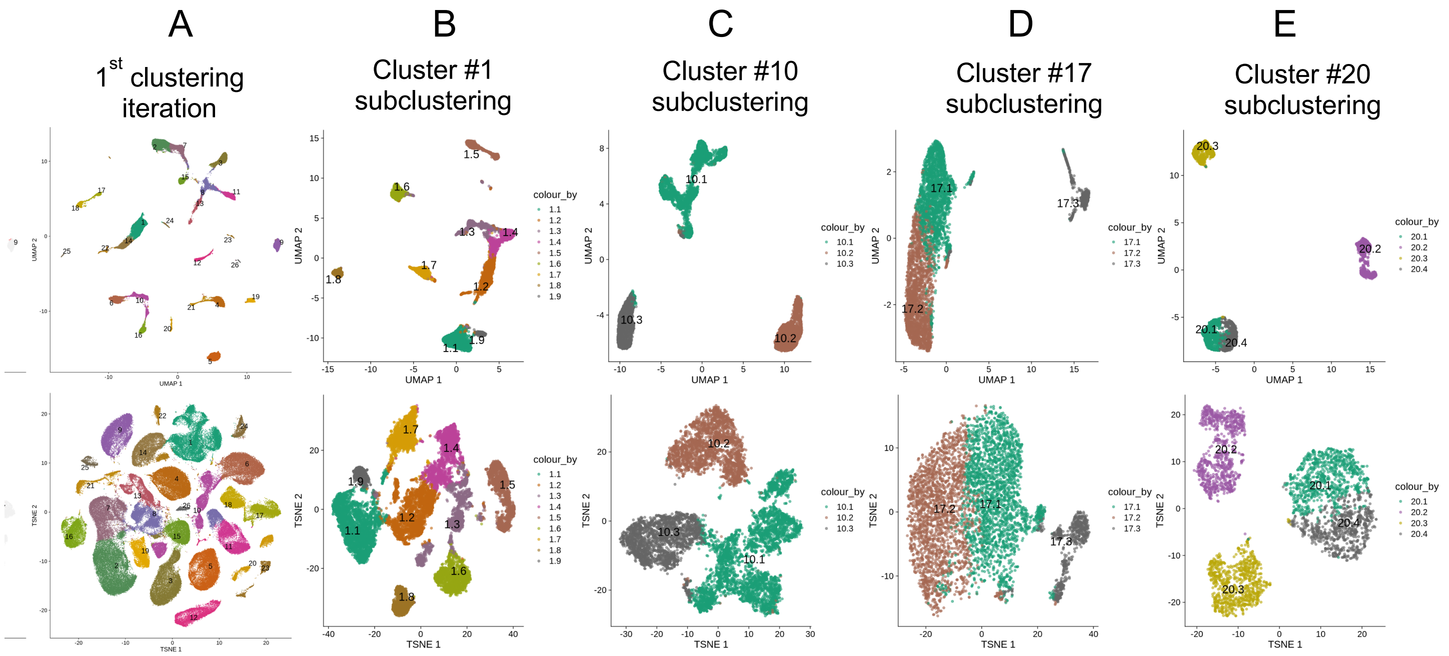


**Supplementary Figure 8 – Sub-clustering results.** UMAP (top) and tSNE (bottom) embeddings are shown. A) The initial set of 26 clusters identified by Leiden community detection are shown. The results of a second round of sub-clustering using Leiden community detection for Clusters 1 (B), 10 (C), 17 (D), and 20 (E) are shown. Thirteen of the original 26 clusters were selected for sub-clustering, resulting in a final total of 60 iterative clusters for further analysis.
